# Nutrient competition predicts gut microbiome restructuring under drug perturbations

**DOI:** 10.1101/2024.08.06.606863

**Authors:** Handuo Shi, Daniel P. Newton, Taylor H. Nguyen, Sylvie Estrela, Juan Sanchez, Michael Tu, Po-Yi Ho, Qinglin Zeng, Brian DeFelice, Justin Sonnenburg, Kerwyn Casey Huang

## Abstract

Human gut commensal bacteria are routinely exposed to various stresses, including therapeutic drugs, and collateral effects are difficult to predict. To systematically interrogate community-level effects of drug perturbations, we screened stool-derived *in vitro* communities with 707 clinically relevant small molecules. Across ∼5,000 community–drug interaction conditions, compositional and metabolomic responses were predictably impacted by nutrient competition, with certain species exhibiting improved growth due to adverse impacts on competitors. Changes to community composition were generally reversed by reseeding with the original community, although occasionally species promotion was long-lasting, due to higher-order interactions, even when the competitor was reseeded. Despite strong selection pressures, emergence of resistance within communities was infrequent. Finally, while qualitative species responses to drug perturbations were conserved across community contexts, nutrient competition quantitatively affected their abundances, consistent with predictions of consumer-resource models. Our study reveals that quantitative understanding of the interaction landscape, particularly nutrient competition, can be used to anticipate and potentially mitigate side effects of drug treatment on the gut microbiota.

## INTRODUCTION

Microbial communities play pivotal roles across natural, industrial, and clinical settings. In recent years, *in vitro* culturing has emerged as a powerful tool for interrogating the complex behaviors exhibited by these communities, such as nutrient competition^1,2^, cross-feeding^3^, and responses to external supplements^4^ and drugs^5,6^. These communities maintain many ecological interactions among the members, offering insights relevant to their *in vivo* counterparts, such as the response to antibiotics^7^. Moreover, the ability to precisely control the environment enables mechanistic investigations, and multi-condition screens can be carried out with throughput orders of magnitude higher than *in vivo*.

Many therapeutics, particularly antibiotics, can have unintended collateral effects vis modifying the growth and composition of the human gut microbiome^8–10^. Moreover, numerous non-antibiotic drugs have been revealed to have direct inhibitory effects on human gut commensals^11–14^. However, due to the myriad interactions in bacterial communities, it remains unclear whether drug sensitivity in isolation translates to inhibition within a community, and how such sensitivity cascades through an interconnected ecosystem. Nutrient competition among community members has been shown to cause competitive release, in which rare opportunistic pathogens expand due to antibiotic inhibition of the commensal bacteria that normally outcompete pathogens^15^. A less susceptible species can be indirectly sensitized when its cross-feeding partner is inhibited^16^. Beyond nutrients, community-driven factors could also modulate susceptibility^17^. For instance, certain species can alter the environmental pH and thus modulate the antibiotic susceptibility of other species^18^. A recent screen of a defined, 32-species synthetic community against 30 drugs showed that while in many cases, species responded to drug perturbations similarly in monoculture and in community, community-specific behaviors involving cross-protection and cross-sensitization can emerge^6^. For example, certain community members can provide general protection through intracellular drug accumulation or degradation^6^. Thus, the outcome of drug perturbation potentially depends on many factors, including species sensitivity and community-level interactions, underscoring the importance of a systematic and quantitative examination of drug–community interactions.

The consequences of collateral damage on the gut microbiota can be severe. Antibiotic usage can shift the human gut microbiome into an alternative, disrupted state^19,20^, potentially because the extinction of certain species opens up nutrient niches for others^2,21^. In some individuals, such disrupted states can persist for several months^22^, and the disruption can facilitate invasion by opportunistic pathogens^19^. Reintroducing commensals via fecal material transplant is often effective in combatting such infections and restoring the gut microbiome to its original state^23,24^. Despite the fundamental importance and clinical relevance of alternative stable microbiota states, systematic studies probing the prevalence and mechanistic determinants of such states are lacking, which limits the development of interventions to reverse a disrupted microbiota.

In this study, we screened 707 small-molecule drugs against multiple stool-derived *in vitro* communities (SICs) from different donors. We identified 141 drugs that significantly affect community growth, composition, and metabolome. The response of individual species to drug treatment was collectively explained by its susceptibility in isolation and the extent of nutrient competition with other community members. Long-term compositional changes resulting from drug treatment were generally due to extinction of susceptible species, which could be reversed by re-introduction of the extinct strains. However, we observed rare cases of strain swapping, in which a phylogenetically related species replaced an original member long after drug removal. We observed few instances of resistance emergence within communities compared to monocultures, highlighting the interplay between resistance selection and community interactions. Finally, we show that nutrient competition alone can cause community-dependent, non-monotonic, yet predictable dose-dependent responses in certain species. Computational modeling based on nutrient competition was largely in agreement with our findings and provided mechanistic insights into the interplay between nutrient competition and drug susceptibility. In summary, our study provided a quantitative framework to predict the responses of microbial communities to chemical disturbances, emphasizing the pivotal role of nutrient competition in community structure and robustness.

## RESULTS

### Chemical screening of a stool-derived *in vitro* community

To investigate the community-level effects of chemical perturbations on gut bacteria, we screened a stool-derived *in vitro* community (SIC) against the National Center for Advancing Translational Sciences (NCATS) clinical collection library. The NCATS library consists of 707 clinically relevant small-molecule drugs that target various types of organisms including human cells, bacteria, viruses, etc. (Fig. 1A, Table S1). Of the 479 for which we were able to identify their administration routes^25^, 402 (84%) can be orally administered. Motivated by a previous *in vitro* screen of gut commensal isolates that discovered widespread susceptibility to human drugs dosed at 20 µM concentration^11^, we initially screened the SIC via anaerobic culturing in the presence of 20 µM of each drug in BHI (Methods). This SIC (hereafter, SIC-0) was derived from a germ-free mouse colonized with a human fecal sample^7^, and consists of a phylogenetically diverse range of bacterial amplicon sequencing variants (ASVs, a proxy for species) representing 14 families (Fig. 1B, 1C). BHI was selected based on our previous findings that SIC composition largely mimics that of the stool from which it was derived^7,26^. During the screen, we monitored community growth dynamics using a plate reader (Methods). At the end of 48 h of incubation with each drug, we assessed community composition via 16S rRNA gene amplicon sequencing (Methods) and assayed the extracellular metabolite profile using untargeted metabolomics (Methods, Fig. 1B).

**Figure 1:**
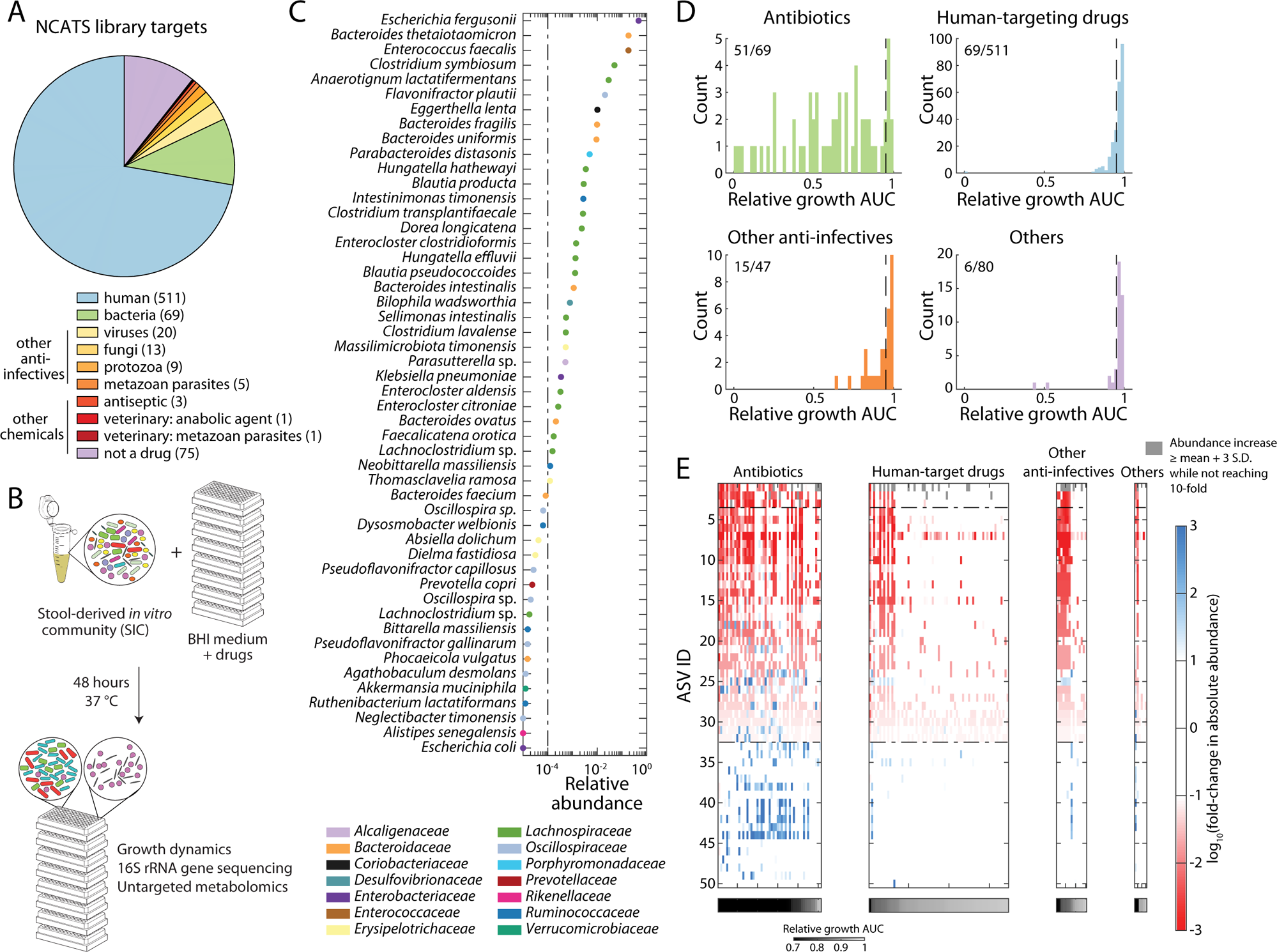
Drug treatment induces large-scale compositional changes in a stool-derived *in vitro* community. A) The NCATS library consists of small-molecule chemicals that target various cell types. Numbers in parentheses denote the total count of chemicals in each category. B) Schematic of the chemical screen. C) Relative abundance of ASVs detected in SIC-0. Each ASV is colored by its corresponding family. Dashed line: limit of detection for a single sample at a typical sequencing depth. SIC-0 was sequenced multiple times, so some ASVs below the limit of detection could be reliably identified. D) While antibiotics generally had large inhibitory effects, many chemicals in other categories also inhibit community growth. Dashed line: threshold for growth inhibition, calculated as the mean–2 standard deviations (S.D.) for control replicates. E) ASV abundance changes in the conditions with growth inhibition identified in (D), with drugs sorted by the extent of growth inhibition in each category. Overall, drug treatment inhibited growth of high-abundance ASVs while allowing low-abundance ASVs to expand. Only fold changes >10-fold or <0.1-fold are shown with colors. ASVs above the top dashed line had initial relative abundance ≥10^−1^, hence could not increase by >10-fold. ASVs below the bottom dashed line had initial abundance ≤10^−4^, hence any decrease in abundance was not detectable. Gray: conditions in which the corresponding ASVs increased in abundance to more than mean+3 S.D. of the control, but not >10-fold. Bottom: growth AUC for each drug condition.

As expected, at 20 µM, most antibiotics (51/69) inhibited community growth (as measured by area under the growth curve (AUC), Fig. 1D, top left). Of the antibiotics that did not substantially inhibit growth, five are typically used specifically to treat the obligate aerobe *Mycobacterium tuberculosis* and likely do not target the anaerobically grown species in SIC-0, and six were beta-lactams from which the community is likely protected via beta-lactamases secreted by certain species^27^. We further analyzed the effects of other drugs by grouping them into human-targeting drugs, anti-infectives targeting viruses, fungi or parasites, and chemicals that are not drugs (Fig. 1A,D). A smaller fraction (14%) of non-antibiotic drugs inhibited growth, and to a lesser average extent than antibiotics (Fig. 1D). Such community-level inhibition is consistent with a previous study^11^ that many human-targeting drugs inhibit the growth of human gut bacterial isolates when grown in isolation. Similar to previous work^11^, we found that antineoplastic agents were enriched in terms of SIC-0 growth inhibition (10/32 inhibited growth compared to 69/511 for all human-targeting drugs, *p* = 0.03, one-tailed Fisher’s exact test).

Variable inhibition of community growth was also accompanied by variable changes in absolute abundance across ASVs. Based on a strategy that we validated in a previous study^1^, we estimated the absolute abundance of each ASV under each condition by scaling its relative abundance (calculated from 16S sequencing) by the final optical density measured at 600 nm (OD_600_) of the treated community and multiplying by the number of colony-forming units of SIC-0 (Methods). Stronger growth inhibition was accompanied by larger inhibition in high-abundance ASVs. For drugs that reduced relative growth AUC to <0.8, the three most abundant ASVs (*Escherichia fergusonii*, *Bacteroides thetaiotaomicron*, and *Enterococcus faecalis*) were all significantly suppressed (*p*<0.001 for all three ASVs, one-tailed Mann–Whitney U test). Drugs that inhibited a broader range of ASVs also generally led to stronger growth inhibition (Fig. S1B, Spearman’s p=−0.65, *p*<10^−10^, two-tailed Student’s *t*-test). Such growth inhibition was also typically accompanied by the expansion in absolute abundance of many low-abundance ASVs, sometimes by >100-fold (Fig. 1E, S1A). These expansions are likely due to the opening of nutrient niches that were originally occupied by strongly inhibited ASVs that were at high abundance^2^, a phenomenon termed “competitive release.” Indeed, isolated strains representing some of these low-abundance ASVs were able to grow to high yield as monocultures (Fig. S1C). Interestingly, drugs with intermediate growth inhibition were associated with the largest number of ASVs that expanded in absolute abundance, presumably because these drugs selectively inhibited high-abundance species while still allowing many other species to grow (Fig. S1B). We further found that regardless of their initial abundances or taxonomies, all ASVs displayed large abundance ranges in response to drug treatments (Supplemental Text, Fig. S1D,E). Thus, our screen generated a diverse range of communities via chemical perturbations.

### Degree of growth inhibition was correlated with changes to both community composition and metabolome

Given the qualitative changes described above, we next explored whether growth was a quantitative predictor of the overall compositional change of the community. Species richness (alpha diversity) did not systematically correlate with our AUC-based growth inhibition metric (Fig. S2A), suggesting that microbiota diversity may be only a crude proxy for microbiota disruption. However, compositional changes as measured via weighted UniFrac distance to the vehicle (DMSO)-treated control were strongly correlated with relative growth (Fig. 2A, Spearman’s p=−0.28, *p*<10^−10^, two-tailed Student’s *t*-test). Similar correlations were observed with other beta diversity metrics (Fig. S2B-D). We did not observe drug conditions that altered community composition without impairing growth. Two drugs (5-nonyloxytryptamine and minocycline) strongly inhibited growth (relative AUC<0.01) but largely did not affect composition, presumably because the inhibition was so strong that the community essentially did not grow during drug treatment and thus composition remained unchanged. Although the various drugs promoted and inhibited distinct bacterial taxa (Fig. 1E, Fig. 2A, bottom), the general correlation between community growth and composition is consistent with competition of overlapping nutrients playing a major role in community assembly^2^. As strains in the community have overlapping nutrient niches, low-abundance strains only have opportunities to expand when their competing high-abundance strains are inhibited.

**Figure 2:**
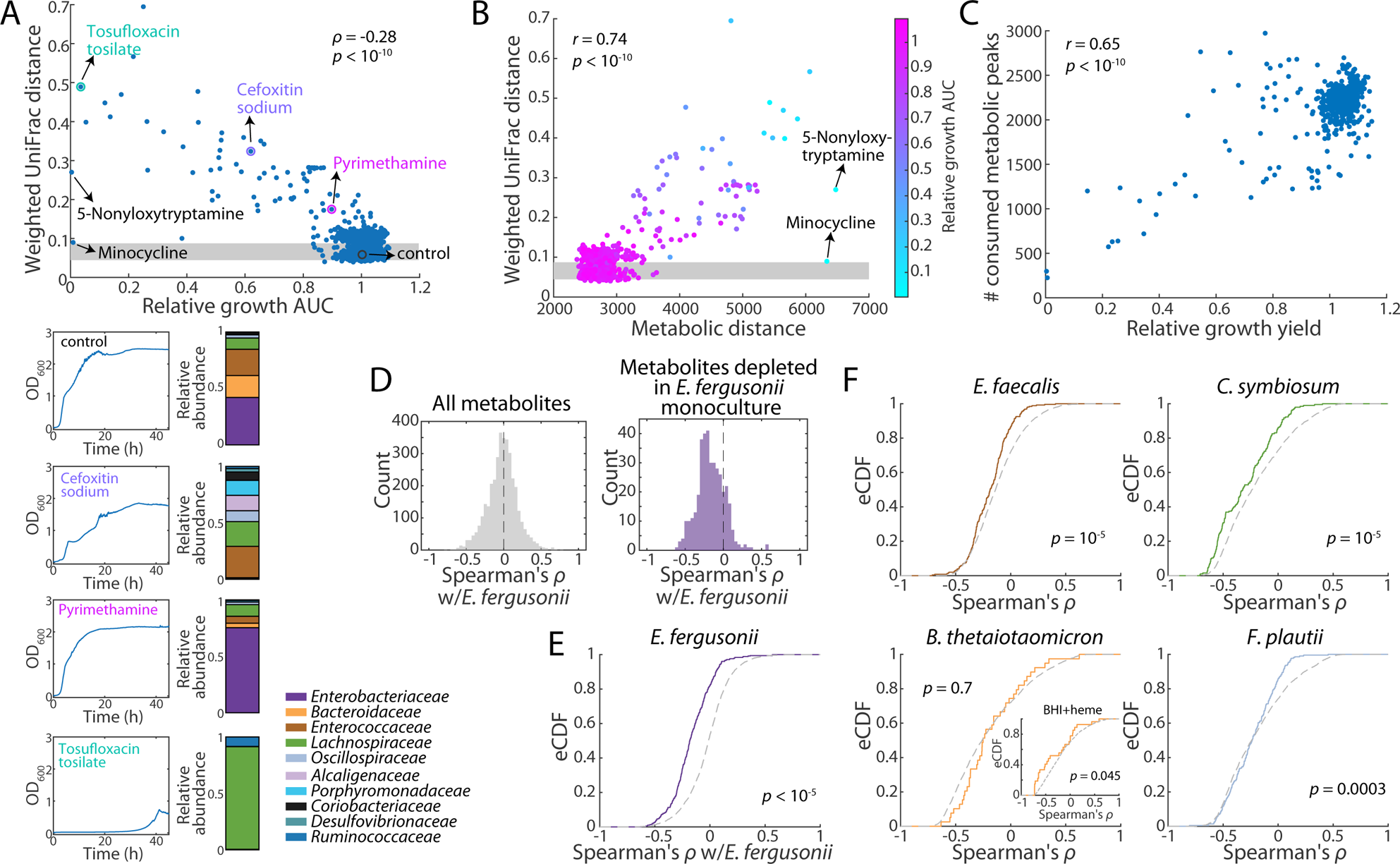
Growth inhibition in a community is correlated with compositional and metabolomic changes. A) Top: for the resulting communities after drug treatment of SIC-0, their relative growth AUC was negatively correlated with the compositional distance (measured by weighted UniFrac) to the vehicle-treated control. Shaded area: mean±1 S.D. of weighted UniFrac distance for vehicle-treated controls. Bottom: the growth curves (left) and composition (right) of selected communities. B) For drug-treated communities, their metabolomic distance to the control was positively correlated with their compositional distance to the control. Communities with large compositional and metabolomic shifts tended to exhibit larger growth defects, as shown by the color of the circles. C) For drug-treated communities, the number of consumed metabolites was positively correlated with growth yield. D) Left: the background distribution of Spearman’s ρ between each metabolic feature and *E. fergusonii* absolute abundance. Right: the distribution of Spearman’s ρ for only the features that were depleted in *E. fergusonii* monoculture. E) Empirical cumulative distribution function (eCDF) of Spearman’s ρ between all metabolite features (gray) and those depleted in monoculture (purple). The metabolites depleted in monoculture were more likely to have negative ρ values (*p*<10^−5^, one-tailed KS-test), suggesting that *E. fergusonii* consumes a similar set of metabolites in the communities compared to monoculture. F) eCDF of Spearman’s ρ across all metabolites (gray) and those depleted in monoculture (colored by family) for other ASVs present at >1% in SIC-0. There were significant shifts toward negative ρ values for all ASVs except *B. thetaiotaomicron*. Inset: when considering metabolite features depleted by *B. thetaiotaomicron* monoculture growth in BHI supplemented with heme, the eCDF was also shifted to the left.

To test the hypothesis that nutrient competition restructures community composition, we sought to directly identify nutrient niches by quantifying the profiles of metabolite consumption after 48-h treatments from untargeted metabolomics. We identified the metabolite features that were depleted compared to the media control, and assumed that each depleted feature represents a nutrient in the medium that is consumed by at least one of the members of the community.^2^ We then calculated the distance between metabolomic profiles of the vehicle control and each community (Methods), and found that the metabolomic distance was strongly positively correlated with compositional change (Fig. 2B; Pearson’s *r*=0.74, *p*<10^−10^, two-tailed Student’s *t*-test). 5-nonyloxytryptamine and minocycline were again outliers, presumably due to lack of growth; as expected, the metabolome of these communities was similar to BHI medium. This finding suggests that community composition and metabolite consumption are intrinsically linked. We also observed a positive correlation between the total number of consumed features and growth yield (Fig. 2C; Pearson’s *r*=0.64, *p*<10^−10^, two-tailed Student’s *t*-test), suggesting that drug treatment indeed opened nutrient niches and hampered overall growth of the communities.

Given this correlation between community metabolism and composition, we wondered to what degree each species in the community occupies the same nutrient niche under drug treatment. To evaluate the extent of niche maintenance, we first focused on the most abundant ASV, *E. fergusonii* (*Ef*), as its high abundance and large range of abundance changes across treatments enabled more accurate quantification of its consumed metabolites. For each metabolite feature, we calculated the Spearman’s correlation between metabolite level and *Ef* absolute abundance and found that there was a large spread of correlation coefficients, ρ, across metabolites (Fig. 2D, left). Metabolites exhibiting strongly negative coefficients are likely uniquely consumed by *Ef*, since the degree of metabolite depletion could be predicted from *Ef* abundance alone. We then compared our metabolomics data with a previously published data set of metabolomic profiles of 15 strain isolates^2^ from the same human donor used to generate SIC-0^7^, and identified the metabolites that were depleted by >100-fold in *Ef* monoculture. Compared to the full set of 3599 metabolites, the *Ef*-depleted subset (382/3599, ∼11%) were more likely to exhibit negative ρ across drug-treated communities (Fig. 2D, right). To directly compare the two distributions, we compared their empirical cumulative distribution functions (eCDF), and found a significant shift to negative ρ values in the *Ef*-depleted subset (Fig. 2E). This overlap in consumption between monoculture and community suggests that metabolite consumption by *Ef* is typically independent of the presence of other species or chemical perturbations.

We extended this analysis to all other high-abundance ASVs (relative abundance >0.01 in SIC-0, Fig. 1C) for which we had metabolomic data for isolate monocultures^2^. For *E. faecalis*, *Clostridium symbiosum*, and *Flavonifractor plautii*, the eCDF of the subset of monoculture-consumed metabolites was shifted to negative ρ values compared to all metabolites (Fig. 2F), suggesting that these species also consume a similar set of metabolites regardless of community or chemical setting. Interestingly, *B. thetaiotaomicron* (*Bt*) was the lone outlier that did not exhibit a significant shift in eCDF (Fig. 2F). We hypothesized that this apparent discrepancy could be explained by the poor growth of *Bt* in BHI monoculture, while its growth is boosted in the presence of *Ef*^2^, potentially by the production of iron-binding siderophores. To test this hypothesis, we analyzed metabolomics data from *Bt* monocultures grown in BHI supplemented with heme. Heme is an iron-containing compound that supports substantially more *Bt* growth than BHI alone. In this case, the eCDF of monoculture-consumed metabolites shifted significantly to the left (Fig. 2F), suggesting that *Bt* also maintains part of its niche in a community and across drug treatments. Moreover, this finding regarding *Bt* metabolism highlights the importance of identifying non-competitive interactions and/or environmental variables that impact species growth in a community and thereby mimic the community growth environment as closely as possible when comparing monoculture and community growth. Taken together, our metabolomics data reveal that although community members compete for nutrients, high-abundance species still have access to distinct and conserved nutrient niches regardless of community context or drug perturbations.

### Drug susceptibility and nutrient availability collectively alter ASV abundance

We next queried the extent to which drug susceptibility during community growth was directly determined by susceptibility in monoculture. We assayed the susceptibility of 14 isolates from SIC-0 against 43 drugs (Fig. 3A) by measuring the final OD_600_ after growth to saturation (Methods). These 14 isolates account for ∼92% of the abundance in SIC-0 and represent six families. Twelve of the isolates are representatives of species that are among the top 20 most abundant in SIC-0 (Fig. 1C), and we further included *Klebsiella pneumoniae* (*Kp*) and *Akkermansia muciniphila*, since both exhibited large changes in abundance in selected conditions (Fig. S1D). Since some isolates grew poorly in BHI medium as monocultures, we supplemented with heme for the Bacteroidota strains and with mucin for *A. muciniphila* (Methods) to more accurately quantify growth inhibition. The 43 drugs, selected to encompass a range of growth inhibitions, included 30 antibiotics from different classes of mechanism of action, 7 human-targeting drugs, 3 antifungals, 1 antiparasite, 1 antiviral, and 1 antiseptic (Table S2). Similar to a previous study^28^, we did not observe any strong trends between phylogenetic relatedness and drug responses in these isolates (Fig. S3A).

**Figure 3:**
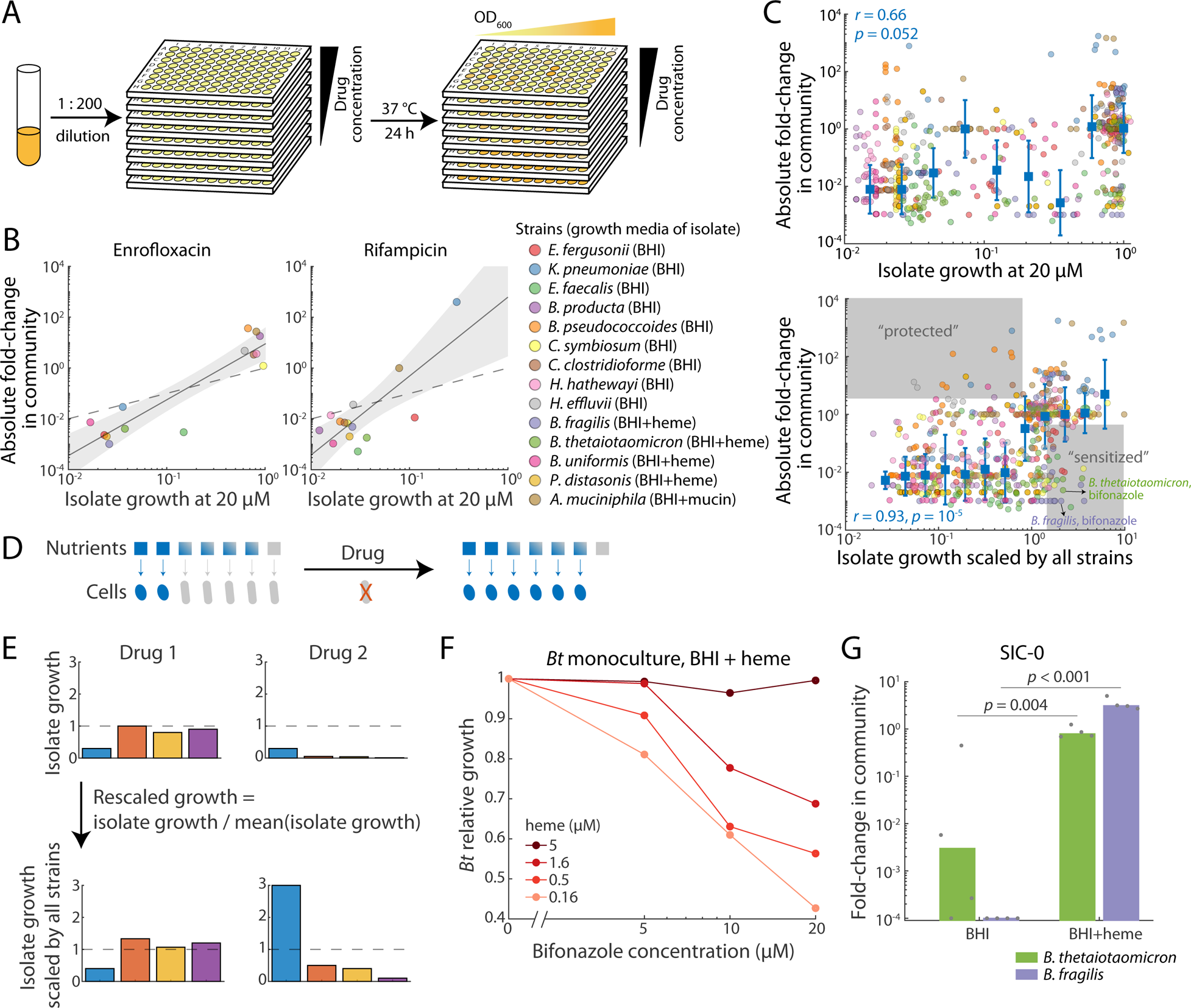
Growth inhibition in a community is collectively explained by isolate susceptibility and nutrient competition. A) Schematic of drug susceptibility quantification for strain isolates. B) After drug treatment, isolate growth in monoculture was largely correlated with fold change in the community. For enrofloxacin treatment (left), most data were close to the line *y*=*x* (dashed line), signifying quantitatively similar susceptibility. However, for rifampicin treatment, although *K. pneumoniae* growth was inhibited by ∼70% in monoculture, since all other species were more impacted, *K. pneumoniae* expanded by >100-fold in the community, presumably due to lack of competition. Dashed lines: *y*=*x*; solid lines and shaded areas: best linear fits and 95% confidence intervals. C) Top: Isolate susceptibility was not correlated with absolute fold change in the community. Bottom: When rescaled to account for the variable growth across strains during treatment in monocultures, isolate susceptibility was correlated with absolute fold change in the community. Blue squares and error bars represent mean±1 S.D. of binned data. D) Simplified schematic of nutrient competition resulting in apparent protection. The blue and gray cells each exclusively consume some nutrients (blue and gray squares, respectively) and also share some nutrients (squares with blue/gray gradients). Without drug treatment, the gray species outcompetes the blue species by consuming all shared nutrients. However, for drug treatments that selectively inhibit the gray species, the blue species has access to all shared nutrients and increases in abundance. E) Schematic of growth rescaling strategy in a simplified scenario with four isolates. The blue isolate has the same susceptibility to drugs 1 and 2, but due to differential susceptibilities of the other strains, its rescaled growth is very different, reflecting the potentially large role of nutrient competition. F) *B. thetaiotaomicron* was sensitized to bifonazole with low levels of external heme. G) Heme supplementation rescued *Bacteroides* species during bifonazole treatment. Bars are averages of *n*=4 biological replicates (individual circles). *p*-values are from two-tailed Student’s *t*-tests.

Our baseline hypothesis was that the relative growth of a monoculture in the presence of a drug should be correlated with the absolute abundance of the species in SIC-0 during treatment. This hypothesis was largely borne out for some drugs (Fig. 3B), but not others. For the fluoroquinolone antibiotic enrofloxacin, the data points were somewhat close to the *y*=*x* line (Fig. 3B, left). In other conditions treatments such as with the ansamycin antibiotic rifampicin, the data points substantially deviated from *y*=*x* (Fig. 3B, right). In this case, all isolates were substantially inhibited in monocultures. Nonetheless, the absolute abundance of *Kp* in the community increased by >100-fold while its monoculture growth was only ∼30% compared to a vehicle-treated control. Similar protection of otherwise susceptible species in a community was reported in a recent study^6^. Although many community-based factors could underlie such boost in *Kp* growth, a simple explanation is that other strains, such as the high-abundance species *E. fergusonii* (*Ef*) that belongs also to the *Enterobacteriaceae* family as *Kp*, compete for the same resources as *Kp*. As these competitors were more strongly inhibited, *Kp* was able to expand in absolute numbers in a community despite its lower growth potential in isolation. We observed similar instances of such competitive release of less susceptible species with other species and drug conditions (Fig. 1E, S1A). Overall, we found that monoculture growth and abundance change in community were poorly correlated across the species and drug conditions tested (Fig. 3C, top), consistent with previous findings in a defined bacterial community.^6^

We sought to test whether nutrient competition is the main factor confounding the correlation between monoculture and community response to drugs. Given that many species in SIC-0 have overlapping nutrient niches,^2^ we postulate that rather than the *absolute* growth inhibition in monoculture, the *relative* growth inhibition with regard to other community members matters more. A species with less *relative* growth inhibition can have growth advantage due to more accessible nutrients freed up from other competing species. For example, in a hypothetic case where two species compete for nutrients, drug treatment killing the dominating species could allow the other species to expand since it has access to more nutrients (Fig. 3D). To approximately account for the effects of nutrient competition, for each drug condition, we rescaled the growth of each isolate by the average growth of all isolates, as a proxy for *relative* growth (Fig. 3E). Thus, a species having the same *absolute* growth under two drug conditions (blue species, Fig. 3E) can have vastly different scaled growth, depending on the susceptibilities of other species. Across all species and drug conditions tested, the rescaled growth was correlated with the corresponding fold-changes in community after treatment (Fig. 3C, bottom). Thus, although the rescaling does not account for the details of niche overlaps across species or other community-level interactions such as cross-feeding, it suggests that inter-strain competition greatly shapes the community response. While we cannot rule out that competition could come from various sources such as toxin production, the correlation across all species tested (Fig. 3C, bottom) suggests nutrient competition to be the most likely factor as it does not depend on specific species. Therefore, this finding indicates that the abundance of a strain is tuned both by its drug susceptibility and competition with other strains in the SIC, and it also provides further support that nutrient competition is the dominant interaction mechanism for this SIC during growth in BHI.

We next asked whether certain strains appeared to be further protected or sensitized (Fig. 3C, bottom, gray regions; Methods) in a community setting beyond the effects of resource competition. Of all 602 strain-drug pairs, 15 (2.5%) were protected and 42 (7.0%) were sensitized. The few cases of potential protection all involved low-abundance species (<1% relative abundance in SIC-0). For 12 of the sensitized conditions, drug treatment enabled the expansion of another ASV for which we lacked monoculture data; hence, the corresponding rescaled growth was likely an over-estimate. Of the remaining sensitized conditions, 11 involved *Bacteroides* species, mostly associated with nitroimidazole chemicals. For example, in the presence of bifonazole, *B. thetaiotaomicron* (*Bt*) and *Bacteroides fragilis* (*Bf*) both grew normally in monoculture, while their abundances dropped by >100-fold during community treatment (Fig. 3C, bottom).

Across bifonazole concentrations, we found that *Bt* was consistently sensitized in a community treated with ≥20 µM bifonazole (Fig. S3B,C). However, we noted that one difference between the *Bt* monoculture and community experiments was that the medium for *Bt* monocultures was supplemented with 5 µM heme to promote growth and acquire reliable OD_600_ readings (Methods). For experiments with SIC-0, no additional heme was needed, likely due to the production of heme by species such as *Ef*^2,29^. High concentrations of heme have been reported to decrease metronidazole susceptibility in other anaerobic bacteria^30,31^. Hence, we hypothesized that high levels of external heme in monoculture protects *Bt* from bifonazole treatment. Indeed, *Bt* was more susceptible to bifonazole in monoculture at lower heme concentrations. While *Bt* monoculture growth was not affected by 20 µM bifonazole treatment when the medium was supplemented with 5 µM heme, growth was inhibited by ∼60% with 0.16 µM heme supplementation (Fig. 3F), even though 0.16 µM heme still supported some *Bt* growth in BHI (Fig. S3D). When *Bt* isolate susceptibility with 0.16 µM heme was used to recalculate rescaled growth, *Bt* was no longer be classified as a “sensitized” outlier. Indeed, we grew SIC-0 in BHI supplemented with 5 µM heme, and found that both *Bt* and *Bf* no longer appeared sensitized to bifonazole treatment (Fig. 3G). Taken together, while *Ef* likely cross-feeds *Bacteroides* with heme under normal growth conditions, the amount of secreted heme is not sufficient to support *Bacteroides* growth in the presence of bifonazole. Compounds like heme that affect both growth and drug sensitivity, although rare in our screen, highlight the interplay between media composition, cross-feeding, and community growth.

### Extinction restructures post-recovery community composition

In humans, a short interval of antibiotic treatment often causes the gut microbiome to shift to an alternative state with lower diversity^20^. In certain individuals, such alternative states can persist for months before species from the environment recolonize^22^. To assess the robustness of communities during recovery from drug perturbations, we performed an additional passage after treatment of SIC-0 (Fig. 1B) by diluting the resulting communities 1:200 into fresh BHI without drug (Fig. S4A). We found that the additional passage led to partial recovery in growth, composition, and metabolomics (Supplemental Text, Fig. S4B).

We next asked whether the lack of full recovery was because perturbed communities require multiple passages to return to equilibrium^7^, or due to permanent strain extinctions. We selected 23 drugs with a range of growth inhibitions, treated SIC-0 with 20 µM as before, and recovered the resulting communities in fresh BHI for six passages (R1–R6, Fig. 4A). After the dilution, up to ∼0.1 µM of drug was carried over into R1. In a previous study quantifying the MIC of 63 antibiotics against five strains, there were only 13 instances (4%) with MIC close to or less than 0.1 µM^32^. Thus, in most cases, drug carryover should not affect community growth even in R1, let alone R2-6. To interrogate the effects of extinctions, we also carried out a parallel set of six recovery passages in which untreated SIC-0 was seeded into treated communities at 1% relative abundance (v/v) directly after treatment (Fig. 4A), mimicking the effect of reseeding susceptible species from environmental reservoirs to the host gut after antibiotic perturbations^33^. For an ASV present at ∼10^−3^ relative abundance in SIC-0, this strategy seeds ∼10 cells into the new passage. Each condition was carried out in duplicate to identify any stochasticity in the outcomes. Overall, we observed high reproducibility across the replicates regardless of seeding conditions (Supplemental Text, Fig. S4C-E).

**Figure 4:**
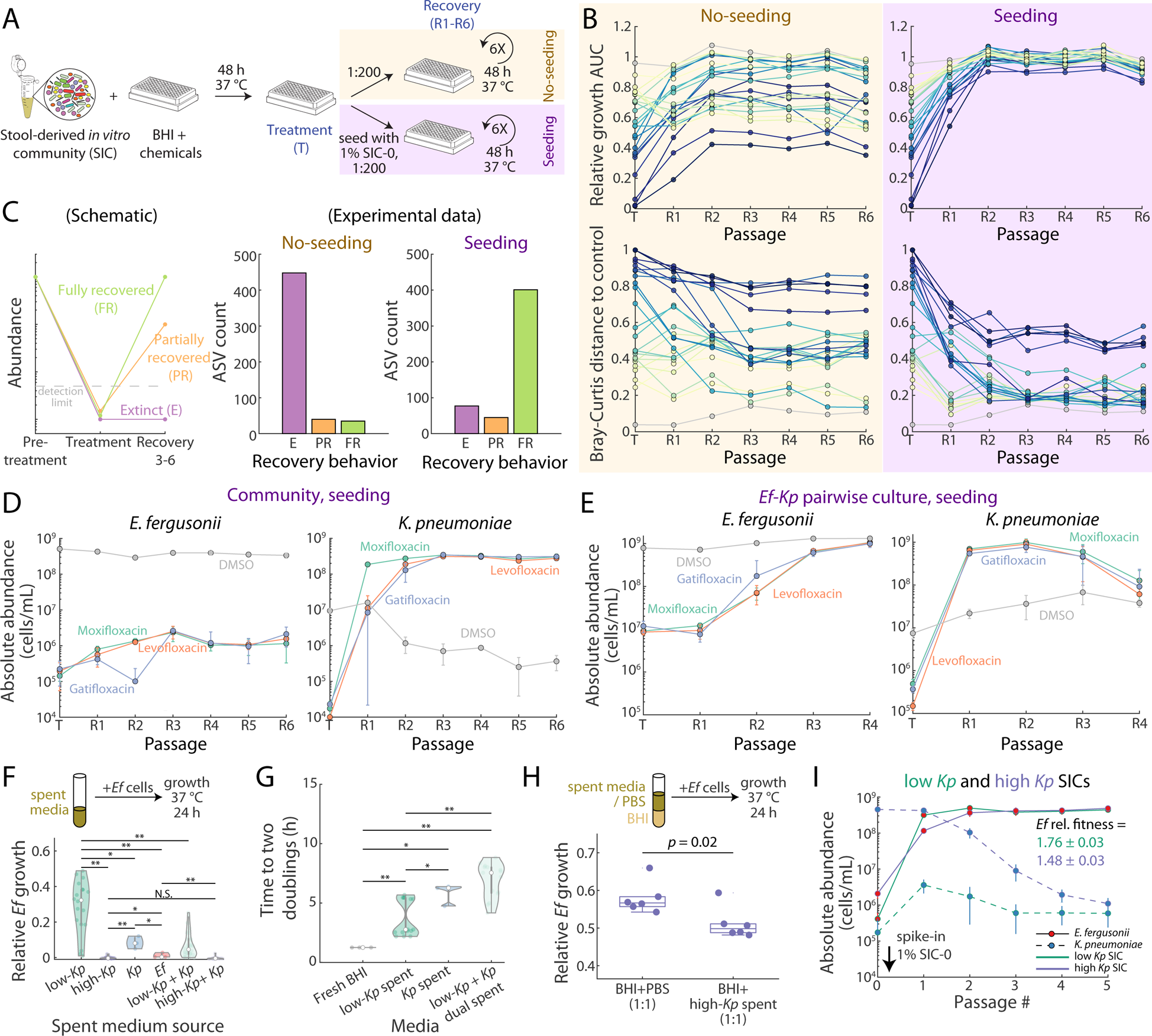
Recovery from drug treatment is hampered by extinction. A) Schematic of multiple recovery passages with or without seeding with SIC-0. B) Both growth dynamics (top) and composition (bottom) required 2–3 recovery passages to equilibrate. With seeding, communities exhibited greater recovery of growth and compositions more similar to the vehicle-treated control than without seeding. Colors represent different drugs, and data are the mean of *n*=2 replicates. C) Left: schematic of definition of ASVs as extinct (E), partially recovered (PR), or fully recovered (FR). Dashed line: limit of detection. Middle: without seeding, most ASVs not detected during treatment eventually went extinct. Right: with seeding, many more ASVs fully recovered compared with no seeding. D) Even with seeding, *E. fergusonii* (*Ef*) did not fully recover after fluoroquinolone treatment (left), and its niche was filled by *K. pneumoniae* (*Kp*, right). Data points and error bars are mean±1 S.D. across *n*=2 replicates. E) In an *Ef*-*Kp* pairwise co-culture with seeding of the pairwise mixture, fluoroquinolone treatment temporarily promoted *Kp* growth, but *Ef* re-emerged after 2–3 recovery passages. Data points and error bars are mean±1 S.D. across *n*=4 replicates. F) Top: *Ef* growth was assayed in spent media from various cultures. Bottom: While spent media from high-*Kp* communities did not allow for *Ef* growth, those from low-*Kp* communities allowed partial *Ef* growth. In dually spent media after *Kp* growth in low-*Kp* spent media, *Ef* growth was further limited. *: *p*<0.05, **: *p*<0.01, N.S.: not significant, one-tailed Mann–Whitney U tests. G) Although spent media from low-*Kp* communities, *Kp* alone, and low-*Kp* communities + *Kp* dually spent media allowed for some *Ef* growth, the time required for *Ef* to double twice in the spent media was significantly longer than in fresh BHI medium, suggesting that the leftover nutrients are less preferable to *Ef*. *: *p*<0.05, **: *p*<0.01, one-tailed Mann–Whitney U tests. H) In BHI supplemented with the spent media from high-*Kp* communities, *Ef* growth was inhibited compared to supplementation with PBS. *n*=6, one-tailed Mann–Whitney U test. I) For the communities with low *Ef*, spiking in SIC-0 enabled *Ef* to quickly take over. If *Kp* was not present in the community, *Ef* expanded even faster. Data points and error bars are mean±1 S.D. of *n*=4 replicates.

Regardless of seeding, communities required 2–3 recovery passages to equilibrate after treatment, after which the variation in growth and composition across passages was similar to that of the vehicle control (Fig. 4B). Without seeding, recovery typically remained partial, with many drugs resulting in large-scale, long-lasting growth inhibition and compositional changes (Fig. 4B, left). With seeding, communities exhibited more recovery, with higher growth and more similar composition to the control (Fig. 4B, right). To investigate the promotion of recovery by seeding, we focused on the ASVs that appeared to go extinct during treatment. Across all conditions, there were 523 ASV– drug combinations in which an ASV was present before treatment but not detected at the end of the treatment passage. We classified these apparent extinctions into three categories based on the ASV’s equilibrated abundance after recovery (average of R3– R6): extinct (not detected at recovery), partially recovered (recovery abundance ≤10% of pre-treatment), and fully recovered (recovery abundance >10% of pre-treatment) (Fig. 4C, left). Without seeding, 86% (448/523) of apparent extinctions during treatment remained extinct during recovery (Fig. 4C, middle). The ASVs that fully recovered without seeding were initially present at lower abundances than those that remained extinct (*p*<0.001, two-tailed Mann–Whitney U test). This bias is likely because low-abundance ASVs can appear extinct (below the limit of detection) even when growth inhibition is only mild, while high-abundance ASVs require larger inhibition to appear extinct and hence are less likely to recover after drug treatment. In contrast, with seeding, 77% of apparent extinctions (401/523) fully recovered (Fig. 4C, right), and the pre-treatment abundances of fully recovered ASVs were similar to those of extinct ASVs (*p*=0.25, two-tailed Mann–Whitney U test). The higher number of extinctions in the no-seeding conditions were also accompanied by more instances of blooming, in which an initially undetected ASV increased in abundance during recovery (Supplemental Text, Fig. S4F). Regardless of seeding, for the ASVs that were present prior to and at the end of treatment, their equilibrated abundance was largely the same as pre-treatment (slope=0.93±0.03 without seeding, and 1.03±0.03 with seeding), even though in many instances (92/302) drug treatment temporarily decreased the ASV’s abundance by >10-fold. Taken together, drug treatment generally caused extinction of many ASVs, leading to long-term growth defects and compositional changes, and resupplying the extinct ASVs via seeding alleviated most of these effects.

### Higher-order interactions can drive long-term strain swapping after drug perturbations

To probe mechanisms of long-term compositional changes, we focused on the drugs that caused large compositional shifts even with seeding. Among these were three fluoroquinolones (levofloxacin, gatifloxacin, and moxifloxacin), in which after seeding the dominant species, *E. fergusonii* (*Ef*), was almost entirely replaced by its fellow *Enterobacteriaceae* family member, *K. pneumoniae* (*Kp*) (Fig. 4D), while other ASVs remained at pre-treatment levels. Without seeding, both *Ef* and *Kp* remained low (Fig. S5A). We term such a change between two species without affecting the other community members as a “strain swap”. A similar strain swap occurred with rifampicin treatment, in which *B. thetaiotaomicron* was largely replaced by another *Bacteroides* species (*B. intestinalis*; Fig. S5B). In the case of *Ef* and *Kp*, in a pairwise co-culture *Kp* had sufficiently higher fitness that it became dominant during fluoroquinolone treatment, but the strain swap was only transient, lasting for 2-3 recovery passages (Fig. 4E, S5C,D, Supplemental Text). In contrast, for the six seeded fluoroquinolone-treated communities (three fluoroquinolones, two replicates each), we continued to 13 recovery passages and found that *Ef* remained low in four of them (Fig. S5E). Similarly, *Ef* remained low in four of the six unseeded fluoroquinolone-treated communities, and we were not able to identify any specific ASVs in these communities that could explain the long-term suppression of *Ef* (Supplemental Text, Fig. S5F-H).

Since no single ASV could readily explain the inhibition of *Ef* in these communities, we asked whether the rest of the community members are able to collectively outcompete *Ef*. We refer to the four unseeded communities that did not exhibit *Ef* recovery as low-*Kp* communities (Fig. S5G), and the four seeded SICs that did not exhibit *Ef* recovery as high-*Kp* communities (Fig. S5E). We collected spent media from saturated low-*Kp* and high-*Kp* communities and assayed *Ef* growth in each (Fig. 4F, top). Spent media from high-*Kp* communities did not allow for *Ef* growth, while spent media from low-*Kp* communities still allowed 30% of *Ef* growth compared to fresh BHI (Fig. 4F, bottom). Thus, the low-*Kp* community still had available nutrients for *Ef* to grow. However, the time required for *Ef* to increase its biomass by four-fold after inoculation substantially increased in the low-*Kp* spent media (Fig. 4G), suggesting that the leftover nutrients were less favorable for *Ef* growth. Thus, low-*Ef* communities restricted *Ef* growth by collectively occupying many of the *Ef* nutrient niches, especially its preferred nutrients that enabled its fast growth and expansion. *Ef* growth yield and initial growth rate were further reduced in dually spent media from *Kp* grown in the spent media of the low-*Kp* communities (low-*Kp* + *Kp* spent), compared to spent media from low-*Kp* communities, suggesting that *Kp* is a strong competitor of *Ef*. Since *Ef* did not grow in any spent media from a high-*Kp* community, we further tested whether high-*Kp* spent media exerts inhibitory effects on *Ef* growth in a mixture of fresh BHI with an equal volume of PBS or high-*Kp* spent media (Fig. 4H, top). Indeed, the high-*Kp* spent media reduced *Ef* growth compared to PBS (Fig. 4H, bottom). Thus, *Ef* abundance is kept low both by nutrient competition, especially the nutrients that support fast *Ef* growth, and by direct growth inhibition through the release of inhibitory compounds.

While passaging the low-*Ef* communities, we noted that when *Ef* reached an abundance threshold ∼5×10^−3^ (corresponding to ∼5,000 cells transferred to fresh medium in each passage), it continued to expand to high abundance thereafter. This threshold was not observed in monocultures (Fig. S5I). We hypothesized that at the observed abundance threshold *Ef* is at sufficient initial abundance to successfully compete for its preferred nutrient and expand. To test this hypothesis, we focused on the cases in which *Ef* did not recover during passages R8–13 (Fig. S5E,G). After passage R8, we mixed the resulting cultures with 1% of SIC-0 (v/v), and passaged two replicates of each SIC five times. The spike-in of SIC-0 transferred ∼5,000 *Ef* cells into the inoculum. In all cases, SIC-0 spike-in to the high-*Kp* communities led to immediate recovery of *Ef* and a concurrent decrease in *Kp* abundance (Fig. 4I). *Ef* expanded more quickly in the low-*Kp* communities, consistent with the observation that high-*Kp* communities compete better with *Ef* for resources (Fig. 4F,G). In contrast, when we mixed SIC-0 with various amounts of *Kp* monoculture, even at a 1:1 ratio, the community resisted *Kp* invasion and returned to a high-*Ef* state after 2 passages (Fig. S5J), further indicating that *Ef* generally outcompetes *Kp* in a community context if at sufficiently high abundance. Thus, in these communities, *Ef* growth appears to be inhibited until *Ef* reaches a critical fraction (∼5×10^−3^), and the presence of *Kp* slows down but does not prohibit its expansion. Expanding above this critical fraction likely allows *Ef* to effectively outcompete other species before they can exert inhibitory effects. To summarize, such multibody dynamics among *Ef*, *Kp*, and other members in the community represent a complex interplay between nutrient competition, inter-species chemical warfare, and priority effects, which can lead to long-lasting compositional changes over >100 generations of growth.

### Resistance selection is rare in SICs

Antibacterial agents naturally drive the selection for resistance^34^, but the rate of emergence and nature of resistant mutants can vary substantially depending on context^35^. In a community, bacteria can acquire resistance through means like horizontal gene transfer^36^. However, due to nutrient competition in a community context, many species remain at low abundances, which limits the rate of evolution^37^ and reduces the chance that a resistant mutant can successfully compete if resistance comes with a fitness tradeoff^38^. We therefore sought to directly assay selection of resistance in communities. After the drug-treated communities reached equilibrium through multiple recovery passages, we re-treated (Fig. 5A) and compared the two treatments to identify ASVs with differential responses.

**Figure 5:**
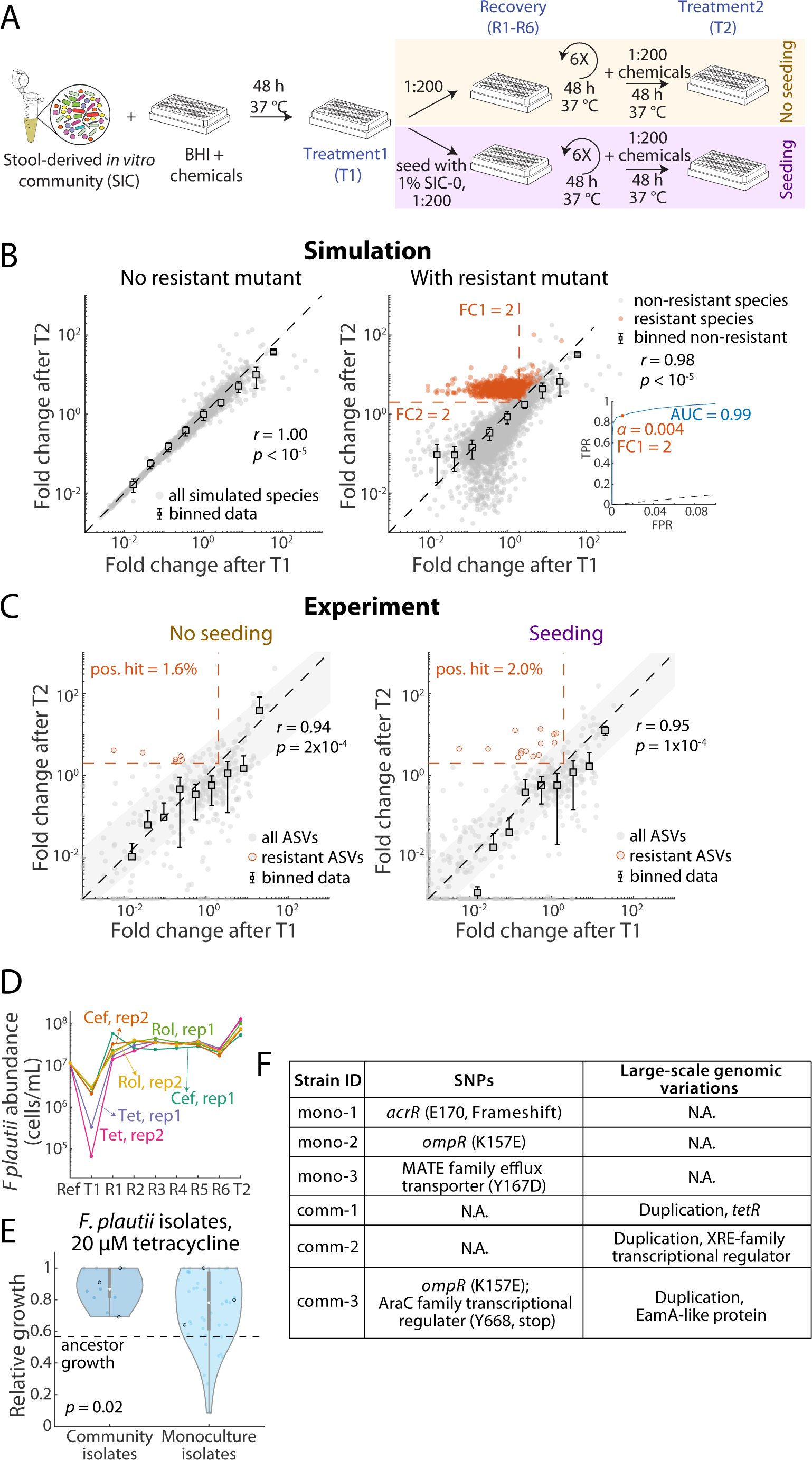
Selection for resistance is rare in communities. A) Schematic of the second round of drug treatment (T2) after recovery to equilibrium. B) Consumer-resource (CR) simulations of communities without (left) or with (right) resistant mutants during the T2 passage. Resistant mutants are colored in orange. Black data points and error bars represent the mean±1 S.D. of binned non-resistant species. Inset: receiver operating characteristic curve for resistant mutants as a function of FC1. C) In the experiments, regardless of seeding condition, most ASVs exhibited similar fold changes across the two treatments, and potential resistance selection (orange outlined dots) was rare. Black data points and error bars represent the mean±1 S.D. of binned data. Shaded areas: 60% confidence interval of linear fit. D) All potential cases of resistance selection without seeding (C, left) involved *F. plautii*. Tet: tetracycline; Cef: cefoxitin; Rol: rolitetracycline. E) Tetracycline-resistant *F. plautii* mutants isolated from the community exhibited higher resistance to tetracycline than those isolated from *F. plautii* monocultures. F) *F. plautii* isolated from tetracycline-treated monocultures or communities had different types of alleles likely to confer tetracycline resistance.

To gain intuition for how resistance would affect the response to drug treatment, we used a consumer-resource (CR) model^39^ to simulate the effects of successive drug treatments during a passaging protocol mimicking our experiments (Methods). Briefly, the model considers 30 species that each consumes a subset of 30 shared resources. Drug treatment was introduced via a random, non-zero death rate for each species. In the absence of resistance, the death rate of a species remains the same for two treatments, and we found that fold changes after the two treatments were strongly correlated and similar in magnitude (Fig. 5B, left), with the small differences due to extinctions in the first treatment that alter the nutrient competition landscape. To model resistance selection, we randomly selected one species and reduced its death rate for the second treatment to be zero. In this case, we observed empirically that the fold change for the resistant species after the second treatment was >2, while all non-resistant species retained approximately the same abundance after the two treatments (Fig. 5B, right). Intuitively, for resistance selection, the fold change during treatment passage T1 should be smaller than that of treatment passage T2. Therefore, we set the cutoff for fold change during passage T1 (FC1) also to be 2. We further estimated the false-discovery rate of such thresholds by varying FC1 and determined from a receiver operating characteristic curve that a threshold of 2 corresponds to a false-discovery rate of 0.004 (Fig. 5B, right).

We used the same strategy to analyze our experimental data and found that, similar to our simulations, ASV fold change across the two treatments was largely the same (Fig. 5C), despite somewhat larger variation presumably due to experimental noise. Moreover, growth AUC across the two treatments was similar for each drug condition (Fig. S5K), consistent with the ASV-level behavior. Thus, we applied the model-inspired thresholds to identify potential cases of resistance. To account for experimental noise, we further excluded cases located within one standard deviation from the line *y*=*x* (Fig. 5C, shaded gray areas). This strategy identified 6 of 376 (1.6%) instances without seeding and 14 of 698 (2.0%) instances with seeding of likely resistance selection (Fig. 5C, circles with red outlines). We then compared these rates of resistance selection with drug-treated monocultures (Fig. 3A). In monocultures, if the final OD_600_ of a strain in the presence of 20 µM of a drug is higher than its final OD_600_ at 10 µM by 1.1-fold, we define the 20 µM condition as having selected resistance. The threshold of 1.1 was chosen because the coefficient of variation across vehicle-treated controls was ∼10%. Across all drugs and species we tested, resistance selection occurred in ∼11% of conditions. Thus, resistance selection is comparatively rare in the context of this community.

Interestingly, all six instances of resistance selection without seeding occurred in *F. plautii* (*Fp*) under tetracycline, cefoxitin, or rolitetracycline treatment (Fig. 5D). Selection for resistant *Fp* mutants was also observed with seeding (Fig. S5L); other instances of resistance selection with seeding involved unique ASVs or occurred in only one replicate of a given treatment. Therefore, we focused on tetracycline treatment to identify the mechanism of resistance in *Fp*. We isolated *Fp* mutants from either tetracycline-treated monocultures or communities (Methods), and obtained 49 isolates from monocultures and 11 from communities. We then assayed their susceptibility to 20 µM tetracycline. As expected, most of these isolates were more resistant to tetracycline than the ancestor (Fig. 5E). However, mutants isolated from the community grew significantly better compared to those from monocultures (*p*=0.02, one-tailed Mann– Whitney U test). To further explore the potential mechanisms of resistance, we performed whole genome sequencing on three isolates from each group (Fig. 5E, circles) as well as the ancestor (Methods). We then searched for SNPs or large-scale genomic structural variations (Methods) that could potentially explain the increased tetracycline resistance in these mutants. In the monoculture isolates (mono-1,2,3), we identified SNPs in the genes *acrR* and *ompR* as well as a gene encoding a MATE family efflux transporter (Fig. 5F). AcrR is a regulator of the AcrAB multidrug efflux pump^40^, OmpR regulates the porin OmpF that controls tetracycline efflux^41^, and certain MATE family efflux transporters can also export tetracycline^42^. In contrast to the monoculture isolates, only one of the three community isolates had SNPs, in *ompR* and an AraC family transcriptional regulator, which is a transcriptional activator related to multi-drug resistance^43^. However, all three community isolates had gene duplications that increased the copy number of potential tetracycline-resistance genes: *tetR*^37^, a xenobiotic response element (XRE)-family transcriptional regulator^44^, and an EamA-like transporter protein^45^. These duplication events likely increased the expression level of these proteins and thereby conferred resistance to tetracycline. Although these genes were present in the ancestral strain, no such duplication events were observed in the monoculture isolates (Fig. 5F). Taken together, these results highlight that *Fp* develops resistance through different mechanisms in monoculture versus in community, potentially because community context biases selection toward resistance mechanisms that are unlikely in monoculture.

### Drug treatment effects are quantitatively adjusted by nutrient competition

To explore whether species respond similarly to drug treatment when embedded in different microbiotas, we screened a random subset (480 of 707) of the drugs against two other SICs, and performed 16S sequencing on selected conditions that exhibited growth inhibition. These two SICs were also derived from mice colonized with the same human fecal sample as SIC-0; one SIC was derived from a mouse switched to a diet deficient in microbiome-accessible carbohydrates (SIC-MD) and the other was derived from a mouse after five days of treatment with the antibiotic ciprofloxacin (SIC-cip)^7,33^. These two perturbations substantially altered microbiota composition in mice, and the composition of the resulting SICs was clearly distinct from SIC-0 while still consisting of a diverse range of ASVs (Fig. S6A). SIC-0, SIC-MD, and SIC-cip consisted of 20, 16, and 20 ASVs present at >10^−3^ relative abundance, respectively. Of these ASVs, SIC-MD shared 12 with SIC-0 (Fig. 6A), and these 12 ASVs accounted for 94% of the total abundance in SIC-0 and 98% in SIC-MD. SIC-cip was more dissimilar from SIC-0 than SIC-MD, sharing only 6 ASVs with SIC-0 (Fig. 6A) that accounted for 68% (SIC-cip) and 24% (SIC-0) of total abundance. Thus, these SICs represent a wide range of initial beta diversities.

**Figure 6:**
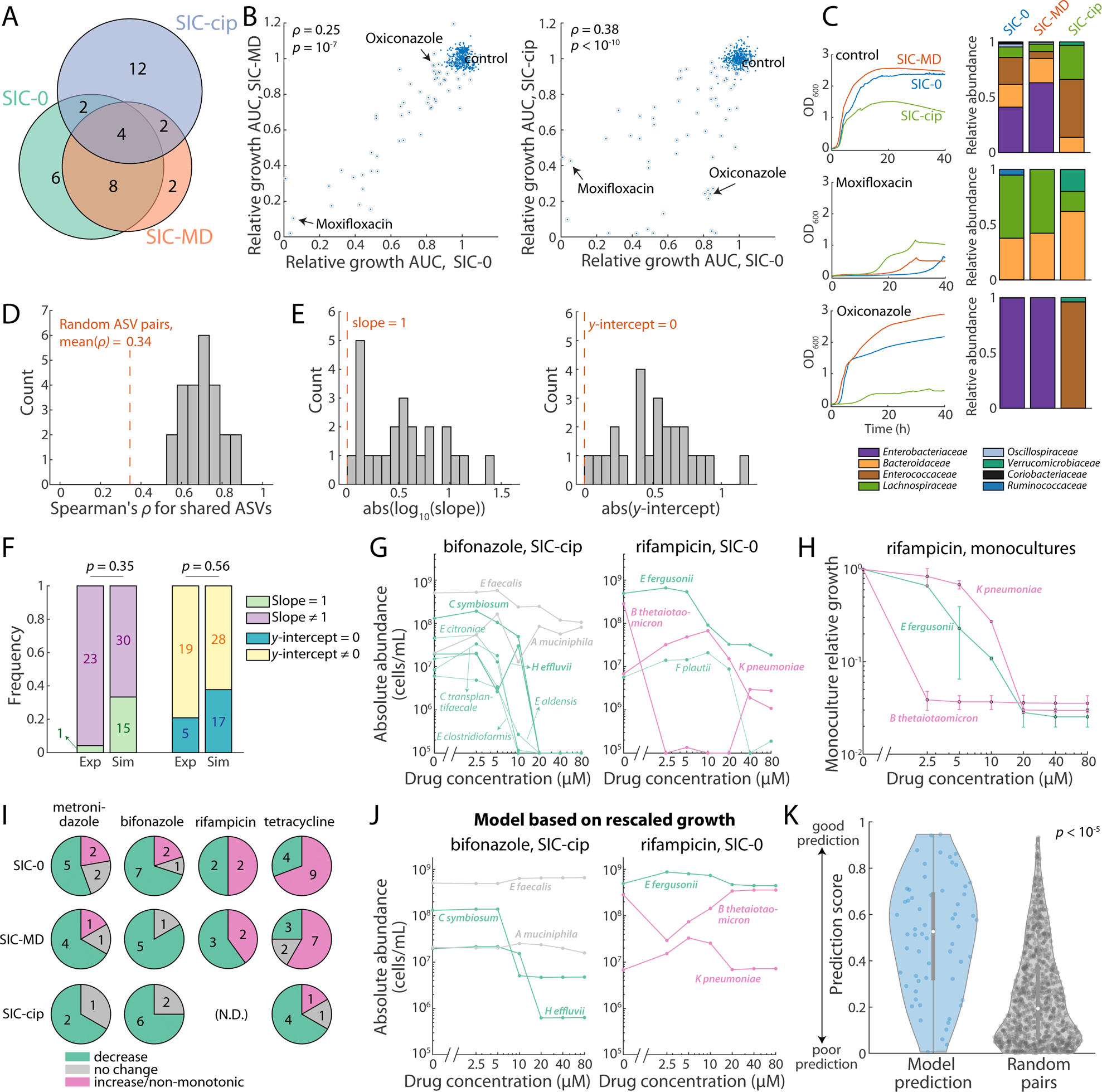
SIC composition quantitatively tunes the effects of drug treatment. A) The three SICs used in our screen have both overlapping and unique ASVs. B) Relative growth AUCs after drug treatment were largely correlated across the three SICs. Samples with 16S amplicon sequencing data are outlined by gray circles. C) Growth curves (left) and family-level compositions (right) of selected drug conditions for the three SICs. During drug treatment, the initial presence or absence of certain ASVs could alter the overall growth response of the community. D) For ASVs shared between SICs, their abundances after drug treatment were correlated. Dashed line: mean Spearman’s ρ for random ASV pairs across SICs, used as a baseline. E) For all ASV pairs in (D), despite strong correlations in abundance, their fold-changes in different SICs generally did not fall along the line *y*=*x*. Slopes of the linear fits deviated from 1 (left, red dashed line) and *y*-intercepts deviated from zero (right, orange dashed line). F) In both experimental data (D,E) and CR simulations, species fold-change was affected by community context. Thus, the majority of ASV pairs exhibited fit slopes deviating from 1 and non-zero *y*-intercepts, defined as the 95% confidence intervals of slopes and *y*-intercepts of the linear fits not containing 1 (for slopes) or 0 (for *y*-intercepts). Exp: experiment; Sim: CR simulations. G) Abundance of high-abundance ASVs across drug concentrations. During bifonazole treatment of SIC-cip (left), all ASVs exhibited monotonic dose responses, while rifampicin treatment of SIC-0 (right) resulted in non-monotonic dynamics for several ASVs. ASVs are colored by their dynamical behavior, and thick lines represent ASVs with isolate susceptibility data. H) Although *B. thetaiotaomicron* and *K. pneumoniae* exhibited non-monotonic dynamics during community treatment with rifampicin, their responses to rifampicin in monocultures were monotonic. I) Across the SICs and drugs tested, increasing/non-monotonic dose responses occurred, and were dependent on community context and drug conditions. Numbers denote ASV counts. N.D.: no data (no growth was observed in SIC-cip with 10 µM or higher rifampicin). J,K) ASV dose responses could be predicted qualitatively (J) when incorporating nutrient competition through monoculture susceptibility and rescaled growth, and (K) predictions were significantly better than baseline (random).

Upon drug treatment, the growth dynamics of SIC-MD and SIC-cip exhibited qualitatively similar behaviors as SIC-0, and the reduction in relative growth AUC was largely correlated across all three SICs (Fig. 6B). Cases in which the extent of growth inhibition differed across SICs were largely due to the inherent compositional differences. For instance, moxifloxacin inhibited growth of SIC-0 and SIC-MD but not of SIC-cip (Fig. 6B,C), likely because SIC-cip contained an *A. muciniphila* ASV (∼3% before treatment, Fig. S6A) and a *Phocaeicola vulgatus* ASV (∼13% before treatment, Fig. S6A) that collectively comprised ∼82% of the community after moxifloxacin treatment; these two ASVs were at <0.01% relative abundance in the other two SICs pre-treatment. For oxiconazole treatment, in SIC-0 and SIC-MD, virtually all ASVs were inhibited except for the dominant ASV, *E. fergusonii* (*Ef*). Since SIC-cip contained very low levels of *Ef* (∼0.07%, Fig. S6A), it experienced much larger growth inhibition in oxiconazole (Fig. 6B,C).

We next asked whether drug treatment led to conserved compositional changes across SICs. For each ASV that was present at >10^−3^ relative abundance in two or more untreated SICs, we calculated their absolute fold change after drug treatment. As representative examples, the abundance changes of *Ef* in SIC-0 and SIC-MD and *E. faecalis* in SIC-0 and SIC-cip were highly correlated (Fig. S6B, Spearman’s ρ=0.81 and 0.56, respectively), and correlations were similarly strong for all overlapping ASVs (Fig. 6D, FDR-adjusted *p*≤10^−6^ for all 24 pairs), suggesting that species generally responded similarly to drug treatments across SICs. We performed a Principal Coordinate Analysis on the treated communities, and found that drug treatment exerted some conserved effects across the SICs (Supplemental Text, Fig. S6C). We further extended our drug screen to eight SICs derived from different human hosts^1^, and again found that growth and compositional changes under drug treatment were largely conserved, with exceptions that were largely due to resistant strains that were present at high abundance in a particular community (Supplemental Text, Fig. S6D-F).

Despite these strong correlations, the ASVs did not simply scale their abundances in response to drug treatment by the same magnitude across SICs. For example, the slope of a linear fit to *Ef* fold change in SIC-0 versus SIC-MD was 0.71±0.10 (Fig. S6C, left). That is, for the same drug perturbation, *Ef* abundance typically changed more in SIC-0 compared to SIC-MD. Such deviation from a slope of 1 is likely caused by community-level interactions. For instance, if a focal species has a close competitor in one community but not the other, then treatments that eliminate its competitor will have less of an inhibitory effect on the focal species due to greater resource availability than in the community without the competitor. For *E. faecalis*, in addition to a slope significantly different from 1, the *y*-intercept of the linear fit was non-zero, as many drugs largely inhibited its abundance in SIC-0 but not in SIC-cip (Fig. S6C, right), presumably due to the absence of the strong competitor *Ef* in SIC-cip (Fig. S6A). Across almost all shared ASVs, the slope of the linear fit was significantly different from 1 (Fig. 6E,F, 23/24 with slope≠1 at 95% confidence interval) and the *y*-axis intercept was non-zero (Fig. 6E,F, 19/24 with *y*-intercept≠0 at 95% confidence interval). We queried whether the quantitative differences in drug responses across communities could emerge simply from nutrient competition using CR models to simulate drug-treated communities (Methods), and calculated the linear fit and correlations for the shared species as above. Even in this simplified scenario in which the only interspecies interaction is nutrient competition, many linear fits had slopes deviating from one (30/45, Fig. 6F) and/or non-zero *y*-intercepts (28/45, Fig. 6F). Thus, even though responses are qualitatively predictable across communities, resource competition within a community frequently quantitatively adjusts each species’s response to drug perturbations.

### Non-intuitive dose responses in communities are related to nutrient competition

Given the large range of susceptibilities across isolates to any particular drug (Fig. 3C), we further tested the response of SIC-0, SIC-MD, and SIC-cip to concentrations increasing two-fold from 2.5 µM to 80 µM of four drugs with a wide range of mechanisms of action. In monoculture, bacterial growth generally monotonically decreases with increasing drug concentration, or stays constant if the strain is resistant. For bifonazole treatment of SIC-cip, eight ASVs were detected in at least three of the six concentrations: the absolute abundances of *E. faecalis* and *A. muciniphila* remained largely unaffected, while the other six ASVs monotonically decreased with increasing drug concentration (Fig. 6G, left). In contrast to such canonical behaviors, the dose-dependence of ASV responses in SIC-0 for rifampicin treatment were more complex. While the growth of two ASVs (*E. fergusonii* and *F. plautii*) decreased monotonically, *B. thetaiotaomicron* (*Bt*) and *K. pneumoniae* (*Kp*) exhibited non-monotonicity: at intermediate rifampicin concentrations, *Bt* was undetectable while *Kp* reached its highest abundance (Fig. 6G, right). In monoculture, both *Bt* and *Kp* growth decreased with increasing rifampicin treatment (Fig. 6H), as expected.

Motivated by these behaviors, we systematically classified the dose response of all ASVs with ≥10^−3^ abundance in at least three concentrations of each drug into three categories: monotonically decreasing (45/83 instances), no change (12/83), or increasing/non-monotonic (26/83). Increasing/non-monotonic ASV-drug combinations were more prominent in SIC-0 and SIC-MD, especially for rifampicin and tetracycline treatment (Fig. 6I). Thus, the interplay of drug inhibition and species–species interactions within a community potentiates nonintuitive dose responses.

Motivated by our observation that nutrient competition and monoculture susceptibility can collectively account for the majority of abundance changes in a community setting (Fig. 3C,D), we wondered whether nutrient competition could also explain the nonintuitive dose responses. Using the monoculture dose response data for the 14 strains we tested (Fig. 3A), we calculated their rescaled growth at each drug concentration as in Fig. 3E and predicted their corresponding fold change. Next, we predicted their abundance in the SIC at each drug concentration by multiplying the predicted fold changes with the species’ absolute abundance in the untreated SIC. For the representative conditions in Fig. 6G, this simplified model based on competition-rescaled growth largely captured the behaviors of all ASVs for which we had monoculture susceptibility data (Fig. 6J), including those with increasing/non-monotonic behavior, although the model tended to overestimate low-abundance ASVs. Such overestimation could be due partly to the limited number of species included in the rescaling, since it is likely that other strains in the community further inhibit the growth of these ASVs. We then evaluated the goodness of fit for our prediction by calculating the normalized overlapping areas between each set of predicted and experimental dose-response curves (Methods). These areas were significantly higher than randomly paired prediction and experimental curves (Fig. 6K, *p*<10^−5^, one-tailed Mann–Whitney U test), suggesting that our model reasonably captures growth behavior in a community context across drug concentrations. Taken together, while other non-competitive interactions could also affect susceptibility in a community context, non-monotonic responses can emerge from straightforward interplay of drug inhibition and nutrient competition.

## DISCUSSION

In this study, we assessed the effects of a wide range of drugs on the growth, composition, and metabolome of diverse communities of gut commensals (Fig. 1B, 4A, 5A, S4A). Notably, community growth was largely associated with changes in community composition and metabolomics profile (Fig. 1E, 2A,B, 6C), suggesting that growth can be used as a rapid, high-throughput assay to filter conditions that impact community composition and metabolite consumption/production. Despite inherent compositional differences across SICs (Fig. 1C, 6A, S6A), ASVs exhibited qualitatively consistent behaviors (Fig. 6C,D,I, S6F), suggesting that our findings may have broad applicability to other synthetic or stool-derived communities of gut microbiota grown in the same medium, particularly if the nutrient competition landscape is largely conserved. *In vivo*, many parameters will likely affect the community response. For example, diet will likely affect the community response by directly modulating nutrient competition within community and changing community composition^33^, or by altering drug pharmacokinetics^46^. In these cases, our *in vitro* data provide a baseline for comparison. Despite the complexity of SICs, drug treatment outcomes were highly reproducible across replicates (Fig. S4C,D), indicating that future screens could expand the chemical space explored by performing fewer replicates. Collectively, our findings regarding growth-based filtering, generality across communities, and reproducibility provide a roadmap for streamlining and increasing the scale of future screens.

Although antibiotics generally caused larger growth inhibition than non-antibiotics, many non-antibiotics also disrupted community structure (Fig. 1D,E). Many drugs from our screen are present in the colon during treatment at approximately 20 µM concentrations^11^, hence the disruptive effects we observed could be relevant in clinical settings. The interplay between nutrient competition and drug response is pivotal in shaping microbial community composition and dynamics. Similar to a previous study with defined bacterial communities^47^, we observed a strong correlation between the compositional and metabolomic shifts due to drug treatment (Fig. 2B). This result is consistent with our finding that bacterial species generally metabolize a similar set of nutrients in monoculture and in a community (Fig. 2E,F), which is an underlying assumption for many studies^2,48^ but rarely experimentally tested. Unsurprisingly, the sensitivities of high-abundance species strongly influenced community responses to drug treatment (Fig. 1E, 6C, S6D,E), in agreement with predictions of consumer-resource (CR) models^39^. The extinction of dominant species when inhibition is sufficiently strong (Fig. 1E, 4D, S5A,B) may involve pressure from competition with other species having overlapping nutrient niches^2^, as evidenced by the expansion of low-abundance species during and after drug exposure (Fig. 1E, S1A, 4D, S4F, S5B). In general, the potential for nonintuitive behaviors to emerge from interactions between nutrient competition and drug inhibition in a community context highlights the utility of CR simulations for mechanistic interrogation of microbial ecology (Fig. 5B, 6F). Due to nutrient competition, many strains appeared to exhibit collateral susceptibility or resistance in SICs compared to in isolation (Fig. 3C, top), similar to the conclusions of a previous study^6^. However, we found that after factoring in nutrient competition by rescaling growth, the susceptibility of isolates largely predicted their drug response behavior in a community context (Fig. 3C, 6G,J). This concordance should enable future studies to focus on outlier cases in which non-competitive interactions such as drug sequestration^6^ or environmental modification^18^ strongly shape the drug response landscape.

Post-drug recovery dynamics are pivotal to understanding microbiome stability and robustness post disturbance. In a previous study involving antibiotic treatment of humanized mice, we measured high levels of residual ciprofloxacin two days after the end of treatment^33^. In the *in vitro* experiments in this study, we found that the residual level of some drugs during the first recovery passage could still inhibit the growth of some species that would otherwise reach high abundance (Fig. S5C); although likely rare, this possibility emphasizes the benefits of multiple recovery passages to enable post-treatment communities to stabilize (Fig. 4B). Seeding with the untreated community (Fig. 4A), which reintroduces extinct species, enhances recovery (Fig. 4B,C), similar to the enhancement of microbiome recovery *in vivo* after fecal microbiota transplantation^24^. Ecological theories have also shown that seeding can lead to functional convergence in microbiome assembly^49^. Our seeding approach (Fig. 4A) supplements even low-abundance ASVs (for an ASV with 10^−3^ relative abundance, ∼10 cells were seeded), so ASV extinctions despite seeding (Fig. 4C) are likely due to residual drugs carried over from dilution of the treated community, and/or to competition from other species. Despite pronounced growth inhibition in many cases (Fig. 1E), most ASVs exhibited similar responses to a second treatment with the same drug (Fig. 5C), suggesting that resistance mutants seldom stabilized in the recovered community. Similarly, decreased resistance selection was observed in a two-species co-culture in rifampicin or ampicillin compared to the corresponding monocultures^50^. Such reduced resistance selection could be due to intense nutrient competition within communities, which tempers the fitness advantages of resistance mutations and suppresses their expansion^51,52^.

Although reintroduction of extinct strains generally improved SIC recovery (Fig. 4B), our multiple observations of strain swapping (Fig. 4D, S5B) highlight one potential manifestation of collateral damage to the microbiota that may be difficult to predict, especially if the emergent strain was undetectable prior to treatment. More generally, as drug interventions inhibit certain strains, they may also enable the expansion of other strains, including pathogens. CR models predict that the inhibited strain should eventually re-emerge after drug inhibition is halted as long as the strain persisted or was reintroduced at a low level. Consistent with this picture, *E. fergusonii* was able to outcompete *K. pneumoniae* in pairwise co-culture regardless of the initial ratio of strain abundances (Fig. 4E). Nevertheless, invasive strains such as *K. pneumoniae* could persist over long time scales as a community recovers from drug treatment if the invader(s) and existing community members collectively outcompete the original strain by exhausting its preferred nutrient niche, and persistence can be accentuated by non-competition-based growth inhibition that may become apparent when the targeted species is present at a very low density (Fig. 4D,F,G). The propensity for strain swapping is likely higher in undefined microbial communities, such as SICs, since they tend to harbor a diverse reservoir of low-abundance species as compared with defined synthetic consortia. Yet, even in our experimental system, such swapping events only occurred occasionally. Thus, microbial behaviors in a community may be predominantly predictable via nutrient interactions, with strain swapping events as the exception rather than the norm. Systematically identifying and understanding cases of strain swapping should provide clinical strategies for curing infections by introducing competitive strains that can outcompete pathogens^53,54^ and for anticipating long-term shifts in microbiome composition^20,22^.

Nonintuitive increasing/non-monotonic dose responses to drug treatment can occur in a community setting (Fig. 6G,I). Such responses are unlikely to be due to global changes like pH modulation or cross-protection via drug uptake or breakdown by resistant species, since these global factors would cause most species to exhibit similar behavior. Instead, we found that nutrient competition alone can qualitatively recapitulate these responses (Fig. 6J). For example, the abundance of a very sensitive species may decrease at low drug concentrations. However, as competitors are inhibited at higher drug concentrations, growth of the sensitive species may increase due to the additional nutrient availability. While obtaining ASV-level dose responses is laborious for community-level screens, computational methods combined with isolate susceptibility data (Fig. 3E, 6J) provide the means to identify potential emergent responses and interpret *in vivo* clinical studies in which chemical concentrations are difficult to accurately quantify.

Moving forward, our approaches and findings provide general insight and principles for the execution and interpretation of high-throughput screens, which are now extending beyond drugs to consider the effects of environmental toxins, dietary compounds, and possibly many other molecules to which the gut microbiome may be exposed. Even with the dramatic increase in scale enabled by *in vitro* cultures such as SICs and synthetic communities and by high-throughput measurement techniques, screening community responses can be laborious and expensive. In this work, systematic analyses of our screens identified factors that can simplify and streamline future endeavors. A key question in microbiome is the extent to which community behaviors can be predicted from isolates and their interactions^3,6^. Among all potential interactions, in the case of gut-derived communities, nutrient competition appears to be a fundamental driver of both community assembly^1,2^ and drug susceptibility (Fig. 3C, 6G,J), providing a baseline hypothesis for responses that highlights specific higher-level interactions that extend beyond this simplified framework (Fig. 3F,G). To accurately account for the effects of nutrient competition, many representative isolates that cover a large fraction of the communities of interest must be isolated (Fig. 3C). Such efforts hold the promise of transforming the complex landscape of chemical–microbe interactions into a means of rationally engineering microbiome composition.

## METHODS

### Chemical screening of SICs

SICs were revived from glycerol stocks by inoculating 5 µL of glycerol stock into 1 mL of fresh BHI medium, and incubated anaerobically for 48 h at 37 °C, which allowed the SICs to reach saturation. 2 mM drug stocks were prepared in dimethylsulfoxide (DMSO). For the initial screen, the resulting cultures were diluted 1:200 into 96-well polystyrene microplates (Greiner Bio-One) containing 200 µL of fresh medium. Two microliters of drug stocks (final concentration 20 µM) or DMSO (for the vehicle-treated control) were added to the mix. The microplates were covered with optical film (Thermo Scientific), with a small hole poked at the edge of each well to allow for gas exchange. The microplate was incubated anaerobically in an Epoch2 plate reader (BioTek) with continuous shaking and OD_600_ readings every 7 min. After 48 h, the cultures were harvested for downstream analyses. For recovery passage 1 with seeding, an aliquot of the resulting 48-h cultures was mixed with saturated SIC-0 at 100:1 (v/v), then diluted 1:200 into fresh medium. For other recovery passages, the 48-h culture was diluted 1:200 into fresh medium.

### Growth curve quantifications

A background OD_600_ value measured from a medium-only well was subtracted from all samples. Afterward, OD_600_ readings were corrected for non-linearity using a standard curve as reported previously^55^.

Relative growth area under the curve (AUC) was calculated by adding all the OD_600_ values from 0 h to 40 h of growth and normalizing to that of the vehicle-treated control. This metric was used because it captures the overall dynamics of the entire growth curve.

### 16S rRNA gene amplicon sequencing and analyses for communities

16S sequencing was performed as previously described^56^. Genomic DNA was extracted from 50–100 µL of cultures using a DNeasy UltraClean 96 Microbial Kit (Qiagen), and the 16S rRNA gene was amplified with 515F/806R primer pairs^57^ using Platinum II HotStart PCR Master Mix (ThermoFisher) with the following thermocycler settings: 94 °C for 3 min, 35 cycles of [94 °C for 45 s, 50 °C for 60 s, and 72 °C for 90 s], then 72 °C for 10 min. The resulting PCR products were pooled at equal volume, concentrated, and cleaned up via agarose gel. Pooled libraries were then sequenced with 250-bp paired-end reads on a MiSeq (Illumina).

Sequencing reads were processed with DADA2^58^. Forward and reverse reads were truncated at 240 and 180 bp, respectively. To account for the multiple 16S genes in certain strains^59^, we curated the ASVs by combining those that differed by 1–2 bp and that exhibited strong linear correlations (FDR-adjusted *p*<0.05) across all drug conditions. Taxonomies of the resulting ASVs were assigned using the SILVA reference database^60^. For ASVs at high relative abundance, their taxonomic annotation was manually curated by blasting the 16S sequence against the NCBI database^61^.

### Estimation of absolute abundances

The absolute abundance of an ASV was estimated by scaling its relative abundance by the final OD_600_ of the community and then multiplying that number by 10^9^. The multiplier 10^9^ corresponds to a measurement of colony-forming units for SIC-0 (∼10^9^ CFUs/mL/OD_600_). Thus, absolute abundances are estimates of total cells per mL of culture.

### Alpha and beta diversity

Shannon diversity was calculated as *H* = −∑*_i_ p_i_* ln *p_i_*, where *p_i_* is the relative abundance of ASV *i*, after rarefying all samples to 5,000 reads. Bray-Curtis distance between samples *A* and *B* was calculated as 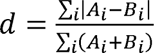, where *A_i_* and *B_i_* are the relative abundances of ASV *i* in communities *A* and *B*. Weighted and unweighted UniFrac distances were calculated using the R function *phyloseq::distance*^62^.

### 16S rRNA gene amplicon sequencing and analyses for isolates

For strain isolates, the full 16S rRNA gene was amplified with 8F/1391R primer pairs (8F: 5′-AGA GTT TGA TCC TGG CTC AG-3′, 1391R: 5′-GAC GGG CGG TGW GTR CA-3′) directly from saturated bacterial cultures following the same PCR protocol as the amplification of genomic DNA from communities. The resulting PCR products were sequenced via Sanger sequencing. Taxonomy was obtained from the full 16S sequences using the NCBI BLAST tool^61^.

### Metabolomics

Bacterial culture supernatants were collected after centrifuging saturated cultures at 4,000*g* and 4 °C for 10 min, and immediately frozen at –80 °C. Before LC–MS/MS analysis, samples were mixed with extraction mix (47.5% acetonitrile, 47.5% methanol, 5% H_2_O, and stable isotope-labelled internal standards), and passed through a 96-well filter microplate with a 0.2-µm polyvinylidene fluoride membrane (Agilent). Samples were injected onto a Waters Acquity UPLC BEH Amide column with an additional Waters Acquity VanGuard BEH Amide pre-column. Spectra were collected using a Thermo Q Exactive HF Hybrid Quadrupole-Orbitrap mass spectrometer in positive and negative mode ionization. Full MS-ddMS2 data were collected. Data were processed using MS-DIAL v. 4.60^63,64^. Alignment retention time and mass tolerance were set to 0.05 min and 0.015 Da, respectively. Protocol details are as previously published^2^.

To calculate distances between metabolomes, we first quantified the changes in area of each MS peak relative to the media control, and defined features as consumed if their concentration was depleted >100-fold and as secreted if their concentration increased >100-fold. The metabolomic distance between samples *A* and *B* was defined as *d*= ∑*_i_*|*A_i_*-*B_i_*|, where *A_i_*(*B_i_*) is –1, 0, or 1 if feature *i* was consumed, unchanged, or secreted, respectively, in sample *A* (*B*).

To identify shared metabolite features from this study and a previous dataset^2^, we first identified the features with identical annotations and *m*/*z* difference within 0.002, and performed a linear fit to correct for retention time (RT) between the two data sets. Unannotated features were matched when the corrected RT difference was within 0.21 min and the *m*/*z* difference was within 0.002. Finally, features exhibiting >10-fold differences across the media controls of the two data sets were filtered out.

### Isolation of *Enterobacteriaceae* strains

SICs were grown anaerobically to saturation in BHI. The resulting cultures were diluted, plated onto LB agar (10 g/L tryptone, 5 g/L yeast extract, 5 g/L NaCl, and 15 g/L agar) supplemented with 10 µg/mL vancomycin, and incubated aerobically for 24 h. The resulting colonies were re-streaked twice onto LB agar with vancomycin, and their identity was verified via 16S rRNA sequencing.

### Drug sensitivity of strain isolates

Glycerol stocks of strain isolates were streaked onto BHI agar plates with 5% horse blood, and for each strain, two independent colonies were picked for downstream experiments. The colonies were grown to saturation in liquid, diluted 1:200 into fresh liquid medium, and arrayed into 384-well microplates. All drugs were dissolved in DMSO at 64 mM and arrayed in 96-well microplates, then added to the 384-well microplates at final concentrations of 320 µM, 160 µM, 80 µM, 40 µM, 20 µM, 10 µM, and 5 µM. The resulting microplates were incubated anaerobically at 37 °C until growth saturated. OD_600_ was measured for the final cultures.

All strains were grown in BHI-based liquid media. To promote growth and obtain reliable OD_600_ readings, the media for all Bacteroidota were supplemented with 5 µM heme, 2 mg/mL NaHCO_3_, and 1 mg/mL L-cysteine, and the media for *A. muciniphila* were supplemented with 5 µg/mL of vitamin K3, 1 µM hemin, 0.2 mg/mL L-tryptophan, 1 mg/mL L-arginine, and 0.5 mg/mL L-cysteine.

In each titration series, if the final OD_600_ at a certain drug concentration was higher than that at a lower concentration, the OD_600_ at the lower concentration was used instead, since the higher OD was likely due to the emergence of resistant mutants^11^. IC_50_ values were calculated as the concentration that corresponded to 50% growth inhibition, as estimated by linear interpolation of the corrected titration series.

To identify potential cross-protection (Fig. 3C, bottom), we used a growth threshold of 0.75, similar to a previous study^6^. The threshold of fold change was set to 3.6, corresponding to a confidence interval of 90% in the control. Similar thresholds (1/0.75 for growth, and 1/3.6 for fold change) were used to identify cross-sensitization.

### Consumer–resource (CR) simulations of communities

Species abundance dynamics during drug treatment were simulated using a CR model as previously described^39^. For the simulations in Fig. 5B, communities were randomly initialized with 30 species with distinct profiles for the consumption of 30 resources. We simulated these communities using the CR model with a constant flow of resources until the composition reached equilibrium. Communities with fewer than 30 co-existing species at equilibrium were discarded, and a total of 1,000 communities with 30 co-existing species were used for downstream simulations. Then, a random death rate sampled from a uniform distribution between 10% and 40% of the dilution rate was imposed for each species to mimic drug treatment, and growth was simulated until equilibrium. The resulting community was allowed to recover to equilibrium with all death rates reset to zero. The community was then exposed to the drug for a second time; for a single species selected to be resistant, the death rate was set to zero while the same death rate was used for the second treatment for all other, non-resistant species. In each round of simulations, species that decreased to relative abundance <10^−4^ were considered extinct and removed from the community.

For the simulations in Fig. 6F, communities were generated with 15 species shared across all communities and 15 unique species. A fixed amount of each resource was supplied, and when all resources were depleted, the resulting abundances were scaled down to mimic a 1:200 dilution, and fresh resources were re-supplied. This passaging was repeated until equilibrium. Three communities with 30 species co-existing at equilibrium were used for downstream simulations. To mimic drug treatment, a random death rate sampled from a uniform distribution was imposed for each species, and the steady-state communities were diluted, supplied with fresh resources, and allowed to grow for a fixed time in the presence of the non-zero death rates. For each community, 200 sets of death rates were used to mimic distinct drug treatments.

### Isolation of resistance mutants

To isolate tetracycline-resistant *F. plautii* mutants from monocultures, a previously isolated *F. plautii* strain from SIC-0^7^ was inoculated from a glycerol stock, and grown overnight. For community, SIC-0 was inoculated from a glycerol stock and grown overnight. The monoculture and community culture were diluted 1:200 into fresh BHI, treated with 20 µM of tetracycline for 48 h, then diluted 1:200 into fresh BHI and grown for another 48 h. The final cultures were diluted and plated onto BHI-blood plates and individual colonies were picked. The *F. plautii* isolate exhibited a flower-like colony morphology, and for the community plates, we biasedly picked colonies with such morphologies and verified their taxonomy via 16S rRNA sequencing.

### Whole genome sequencing of strain isolates

*F. plautii* strains were streaked onto BHI-blood agar plates from glycerol stocks and grown for 48 h. Colonies were scraped, washed in phosphate buffered saline (PBS), resuspended in DNA/RNA shield (Zymo Research), and sent to Plasmidsaurus (Plasmidsaurus Inc., OR, USA) for genomic DNA extraction and sequencing.

Single nucleotide polymorphism (SNP) analyses was performed using SNIPPY v. 4.6.0^65^ with default parameters. Genome alignment was performed with the progressive Mauve algorithm^66^ using Geneious Prime v. 2023.2.1 to identify structural variations.

### Statistics

Statistical tests, *p*-values, and sample sizes are listed in the manuscript for each statistical analysis performed.

## ACKNOWLEDGEMENTS

We thank members of the Huang lab for helpful discussions. We thank the NIH Small Molecule Repository for providing the small-molecule chemical library used in our initial chemical screen. This work was funded by a James S. McDonnell Postdoctoral Fellowship (to H.S.), NSF Award EF-2125383 (to K.C.H.), and NIH Awards R01 AI147023 and RM1 GM135102 (to K.C.H.). K.C.H. is a Chan Zuckerberg Biohub Investigator.

## Notes

### Competing Interest Statement

The authors have declared no competing interest.

## REFERENCES

1 Aranda-Diaz, A. et al. Assembly of gut-derived bacterial communities follows “early-bird” resource utilization dynamics. bioRxiv, 2023.2001. 2013.523996 (2023).

2 Ho, P.-Y., Nguyen, T. H., Sanchez, J. M., DeFelice, B. C. & Huang, K. C. Resource competition predicts assembly of gut bacterial communities *in vitro*. Nature Microbiology, 1–13 (2024).

3 Goldford, J. E. et al. Emergent simplicity in microbial community assembly. Science 361, 469–474 (2018).

4 Celis, A. I., Relman, D. A. & Huang, K. C. The impact of iron and heme availability on the healthy human gut microbiome in vivo and in vitro. Cell Chemical Biology 30, 110–126. e113 (2023).

5 Javdan, B. et al. Personalized mapping of drug metabolism by the human gut microbiome. Cell 181, 1661–1679. e1622 (2020).

6 Garcia-Santamarina, S. et al. Emergence of community behaviors in the gut microbiota upon drug treatment. bioRxiv, 2023.2006. 2013.544832 (2023).

7 Aranda-Diaz, A. et al. Establishment and characterization of stable, diverse, fecal-derived in vitro microbial communities that model the intestinal microbiota. Cell Host Microbe 30, 260–272 e265, doi:10.1016/j.chom.2021.12.008 (2022).

8 Blaser, M. J. Antibiotic use and its consequences for the normal microbiome. Science 352, 544–545 (2016).

9 Zimmermann, M., Patil, K. R., Typas, A. & Maier, L. Towards a mechanistic understanding of reciprocal drug–microbiome interactions. Molecular systems biology 17, e10116 (2021).

10 Maier, L. et al. Unravelling the collateral damage of antibiotics on gut bacteria. Nature 599, 120–124 (2021).

11 Maier, L. et al. Extensive impact of non-antibiotic drugs on human gut bacteria. Nature 555, 623–628, doi:10.1038/nature25979 (2018).

12 Jackson, M. A. et al. Gut microbiota associations with common diseases and prescription medications in a population-based cohort. Nature communications 9, 2655 (2018).

13 Vieira-Silva, S. et al. Statin therapy is associated with lower prevalence of gut microbiota dysbiosis. Nature 581, 310–315 (2020).

14 Vich Vila, A., et al. Impact of commonly used drugs on the composition and metabolic function of the gut microbiota. Nature communications 11, 362 (2020).

15 Varga, J. J. et al. Antibiotics drive expansion of rare pathogens in a chronic infection microbiome model. Msphere 7, e00318–00322 (2022).

16 Adamowicz, E. M., Flynn, J., Hunter, R. C. & Harcombe, W. R. Cross-feeding modulates antibiotic tolerance in bacterial communities. The ISME journal 12, 2723–2735 (2018).

17 Bottery, M. J., Pitchford, J. W. & Friman, V.-P. Ecology and evolution of antimicrobial resistance in bacterial communities. The ISME Journal 15, 939–948 (2021).

18 Aranda-Díaz, A. et al. Bacterial interspecies interactions modulate pH-mediated antibiotic tolerance. Elife 9, e51493 (2020).

19 Costello, E. K., Stagaman, K., Dethlefsen, L., Bohannan, B. J. & Relman, D. A. The application of ecological theory toward an understanding of the human microbiome. Science 336, 1255–1262 (2012).

20 Dethlefsen, L. & Relman, D. A. Incomplete recovery and individualized responses of the human distal gut microbiota to repeated antibiotic perturbation. Proceedings of the National Academy of Sciences 108, 4554–4561 (2011).

21 Spragge, F. et al. Microbiome diversity protects against pathogens by nutrient blocking. Science 382, eadj3502 (2023).

22 Xue, K. S., et al. Prolonged delays in human microbiota transmission after a controlled antibiotic perturbation. bioRxiv (2023).

23 Aas, J., Gessert, C. E. & Bakken, J. S. Recurrent Clostridium difficile colitis: case series involving 18 patients treated with donor stool administered via a nasogastric tube. Clinical infectious diseases 36, 580–585 (2003).

24 Suez, J. et al. Post-antibiotic gut mucosal microbiome reconstitution is impaired by probiotics and improved by autologous FMT. Cell 174, 1406–1423. e1416 (2018).

25 Cai, M.-C. et al. ADReCS: an ontology database for aiding standardization and hierarchical classification of adverse drug reaction terms. Nucleic acids research 43, D907–D913 (2015).

26 Goldman, D. A. et al. Competition for shared resources increases dependence on initial population size during coalescence of gut microbial communities. bioRxiv, 2023.2011. 2029.569120 (2023).

27 Nord, C. E., Lindqvist, L., Olsson-Liljequist, B. & Tuner, K. Beta-lactamases in anaerobic bacteria. Scand J Infect Dis Suppl 46, 57–63 (1985).

28 Smith, T. P. et al. High-throughput characterization of bacterial responses to complex mixtures of chemical pollutants. Nature Microbiology 9, 938–948 (2024).

29 Verweij-van Vught, A., Otto, B., Namavar, F., Sparrius, M. & MacLaren, D. Ability of *Bacteroides* species to obtain iron from iron salts, haem-compounds and transferrin. FEMS microbiology letters 49, 223–228 (1988).

30 Olaitan, A. O. et al. Decoding a cryptic mechanism of metronidazole resistance among globally disseminated fluoroquinolone-resistant *Clostridioides difficile*. Nature Communications 14, 4130 (2023).

31 Paunkov, A., Gutenbrunner, K., Sóki, J. & Leitsch, D. Haemin deprivation renders *Bacteroides fragilis* hypersusceptible to metronidazole and cancels high-level metronidazole resistance. Journal of Antimicrobial Chemotherapy 77, 1027–1031 (2022).

32 Eucast, E. Determination of minimum inhibitory concentrations (MICs) of antibacterial agents by broth dilution. Clin Microbiol Infect 9, ix–xv (2003).

33 Ng, K. M. et al. Recovery of the gut microbiota after antibiotics depends on host diet, community context, and environmental reservoirs. Cell host & microbe 26, 650–665. e654 (2019).

34 Bush, K. et al. Tackling antibiotic resistance. Nature Reviews Microbiology 9, 894–896 (2011).

35 Wistrand-Yuen, E. et al. Evolution of high-level resistance during low-level antibiotic exposure. Nature communications 9, 1599 (2018).

36 Von Wintersdorff, C. J. et al. Dissemination of antimicrobial resistance in microbial ecosystems through horizontal gene transfer. Frontiers in microbiology 7, 173 (2016).

37 Charlesworth, B. Effective population size and patterns of molecular evolution and variation. Nature Reviews Genetics 10, 195–205 (2009).

38 Pinheiro, F., Warsi, O., Andersson, D. I. & Lässig, M. Metabolic fitness landscapes predict the evolution of antibiotic resistance. Nature Ecology & Evolution 5, 677–687 (2021).

39 Newton, D. P., Ho, P.-Y. & Huang, K. C. Modulation of antibiotic effects on microbial communities by resource competition. Nature Communications 14, 2398 (2023).

40 Cuthbertson, L. & Nodwell, J. R. The TetR family of regulators. Microbiology and Molecular Biology Reviews 77, 440–475 (2013).

41 Thanassi, D. G., Suh, G. & Nikaido, H. Role of outer membrane barrier in efflux-mediated tetracycline resistance of Escherichia coli. Journal of bacteriology 177, 998–1007 (1995).

42 McAleese, F. et al. A novel MATE family efflux pump contributes to the reduced susceptibility of laboratory-derived Staphylococcus aureus mutants to tigecycline. Antimicrobial agents and chemotherapy 49, 1865–1871 (2005).

43 Grossman, T. H. Tetracycline antibiotics and resistance. Cold Spring Harbor perspectives in medicine 6, a025387 (2016).

44 Xu, Q. et al. Structure of an MmyB-like regulator from C. aurantiacus, member of a new transcription factor family linked to antibiotic metabolism in actinomycetes. (2012).

45 Jack, D. L., Yang, N. M. & H. Saier Jr, M. The drug/metabolite transporter superfamily. European Journal of Biochemistry 268, 3620–3639 (2001).

46 Harris, R. Z., Jang, G. R. & Tsunoda, S. Dietary effects on drug metabolism and transport. Clinical pharmacokinetics 42, 1071–1088 (2003).

47 Wuyts, S. et al. Consistency across multi-omics layers in a drug-perturbed gut microbial community. Molecular Systems Biology 19, e11525, 10.15252/msb.202311525 (2023).

48 Lopes, W., Amor, D. & Gore, J. Cooperative growth in microbial communities is a driver of multistability. Nature Communications 15, 4709 (2024).

49 Estrela, S. et al. Functional attractors in microbial community assembly. Cell Systems 13, 29–42. e27 (2022).

50 Adamowicz, E. M., Muza, M., Chacón, J. M. & Harcombe, W. R. Cross-feeding modulates the rate and mechanism of antibiotic resistance evolution in a model microbial community of Escherichia coli and Salmonella enterica. PLoS pathogens 16, e1008700 (2020).

51 Fang, P., Elena, A. X., Kunath, M. A., Berendonk, T. U. & Klümper, U. Reduced selection for antibiotic resistance in community context is maintained despite pressure by additional antibiotics. ISME communications 3, 52 (2023).

52 Klümper, U. et al. Selection for antimicrobial resistance is reduced when embedded in a natural microbial community. The ISME journal 13, 2927–2937 (2019).

53 Cabral, V. et al. Gut protective Klebsiella species promotes microbiota recovery and pathobiont clearance while preventing inflammation. bioRxiv, 2023.2011. 2014.566997 (2023).

54 Oliveira, R. A. et al. Klebsiella michiganensis transmission enhances resistance to Enterobacteriaceae gut invasion by nutrition competition. Nature microbiology 5, 630–641 (2020).

55 Atolia, E. et al. Environmental and physiological factors affecting high-throughput measurements of bacterial growth. MBio 11, 10.1128/mbio.01378-01320 (2020).

56 Celis, A. I. et al. Optimization of the 16S rRNA sequencing analysis pipeline for studying *in vitro* communities of gut commensals. Iscience 25 (2022).

57 Walters, W. et al. Improved bacterial 16S rRNA gene (V4 and V4-5) and fungal internal transcribed spacer marker gene primers for microbial community surveys. Msystems 1, e00009–00015 (2016).

58 Callahan, B. J. et al. DADA2: High-resolution sample inference from Illumina amplicon data. Nature methods 13, 581–583 (2016).

59 Větrovský, T. & Baldrian, P. The variability of the 16S rRNA gene in bacterial genomes and its consequences for bacterial community analyses. PloS one 8, e57923 (2013).

60 Quast, C. et al. The SILVA ribosomal RNA gene database project: improved data processing and web-based tools. Nucleic acids research 41, D590–D596 (2012).

61 Johnson, M. et al. NCBI BLAST: a better web interface. Nucleic acids research 36, W5–W9 (2008).

62 McMurdie, P. J. & Holmes, S. phyloseq: an R package for reproducible interactive analysis and graphics of microbiome census data. PloS one 8, e61217 (2013).

63 Tsugawa, H. et al. MS-DIAL: data-independent MS/MS deconvolution for comprehensive metabolome analysis. Nature methods 12, 523–526 (2015).

64 Tsugawa, H. et al. A lipidome atlas in MS-DIAL 4. Nature biotechnology 38, 1159–1163 (2020).

65 Seemann, T. Snippy: Rapid haploid variant calling and core genome alignment, <https://github.com/tseemann/snippy> (2024).

66 Darling, A. C., Mau, B., Blattner, F. R. & Perna, N. T. Mauve: multiple alignment of conserved genomic sequence with rearrangements. Genome research 14, 1394–1403 (2004).

