## Supplemental materials for "Nutrient competition predicts gut microbiome restructuring under drug perturbations"

**Supplemental Information for “Nutrient competition predicts gut microbiome restructuring under drug perturbations”**

**SUPPLEMENTAL TEXT**

**Drug treatment leads to large-scale abundance changes across taxonomies and initial abundances**

We asked whether certain ASVs were generally more susceptible or resistant to drug perturbations by analyzing the range of absolute abundances for each ASV across drugs with inhibitory effects on community growth. We quantified the range of absolute abundance changes as the ratio of maximum to minimum abundance and found that all 50 ASVs varied by >300-fold, with 45 varying >1,000-fold (Fig. S1D). The most abundant ASV, *E. fergusonii*, varied in absolute abundance between <10^6^ and ~10^9^ cells/mL. Even ASVs that were barely detectable in SIC-0 (absolute abundance <10^4^ cells/mL) grew in some conditions to an absolute abundance of ~10^7^ cells/mL. Abundance ranges at the family level exhibited similar variance (Fig. S1E).

**Most drug-treated communities exhibited partial recovery after one recovery passage**

We performed an additional passage after treatment of SIC-0 (Fig. 1B) by diluting the resulting communities 1:200 into fresh BHI without drug (Fig. S4A), and obtained growth, composition, and metabolomics data for the recovery passage. Growth AUC during the recovery passage typically exhibited partial recovery (Fig. S4B): in most cases, communities grew more during recovery than during treatment, although still less than the vehicle-treated control, as signified by a linear fit with slope=0.50 (Fig. S4B, left). Composition (Fig. S4B, middle) and metabolomics data (Fig. S4B, right) also exhibited partial recovery after one passage in fresh BHI, as characterized by linear correlations between the distances of the treated and recovered communities to the pre-treatment community with slopes <1.

**The effects of drug treatment and recovery are reproducible**

In both seeding and no-seeding conditions, most replicates at R6 exhibited highly similar compositions compared to random pairs from different drug treatments (Fig. S4C). In no-seeding conditions, the R6 growth dynamics of replicates were also more similar than those of random pairs; for seeding, all communities at R6 recovered control-like growth dynamics (and hence all replicate pairs grew similarly; Fig. S4D). The fluoroquinolone antibiotic pazufloxacin was an outlier among the no-seeding conditions, in which the two replicates exhibited different compositions (Bray–Curtis distance = 0.72). Closer inspection revealed that the variation was due mainly to the differential abundance of several high-abundance ASVs rather than a global shift in the community (across all ASVs, Spearman’s *ρ*=0.58, *p*<10^–5^, two-tailed Student’s *t*-test; Fig. S4E, left). The pazufloxacin-treated replicate communities with seeding were more similar (Bray–Curtis distance = 0.31); the high-abundance ASVs grew consistently in both replicates. For three of the four ASVs that exhibited large variation across no-seeding replicates (Fig. S4E, left), we had isolated representative strains. In monoculture susceptibility assays, their IC_50_ for pazufloxacin was approximately 20 µM. Typically, bacterial growth in response to drug treatment falls on a logistic curve, with the steepest change occurring around the IC_50_^1^. Thus, these species are more likely to have variable responses across replicates given our screening concentration of 20 µM. In other drugs for which the two replicates had similar composition and growth, all ASV abundances were highly correlated between replicates (Fig. S4E, right). Thus, despite the complexity of SICs, the effects of drug treatment are largely reproducible. The lack of reproducibility in the case of pazufloxacin also highlights that in rare cases, straightforward factors can explain the apparent alternative states of the community.

**Seeding during recovery reduces the persistence of ASVs that bloomed during treatment**

Since certain ASVs expanded in absolute abundance during treatment (Fig. 1E, Fig. S1A), we further asked whether such blooming persisted during recovery. There were 124 instances of ASV-drug combinations in which an ASV that was undetected prior to treatment bloomed during treatment. Without seeding, 44 (35%) of these instances were transient and the ASVs reverted to undetectable levels after recovery, while 80 (65%) persisted at detectable levels throughout all recovery passages (Fig. S4F). Across all conditions, the final abundance of these 80 ASVs was correlated with the total initial abundance of all ASVs that appeared to go extinct (*n*=46 communities, Pearson’s *r*=0.37, *p*=0.008, two-tailed Student’s *t*-test), suggesting that the persistence of blooming species is related to the extinction of their competitors. With seeding, we observed somewhat fewer instances of persistent blooming (68 long-term, 56 transient, *p*=0.03, two-tailed Fisher’s exact test), and those persisted were at much lower abundances than those bloomed without seeding (median abundance of 1×10^7^ cells/mL in unseeded communities vs. 3×10^6^ cells/mL in seeded communities, *p*<10^-5^, one-sided Mann-Whitney U test). The lower frequency and magnitude of persistent blooming with seeding is presumably because seeding prevented some extinctions, and therefore the blooming ASVs faced more competition. Nonetheless, even with seeding, notable persistence (abundance >1×10^7^ cells/mL) still occurred in 14 cases. In summary, even with seeding, drug treatment can result in community reconfiguration long after the drug is removed.

**Strain swapping between *Ef* and *Kp* emerges from community effects**

The swap between *Escherichia fergusonii* (*Ef*) and *Klebsiella pneumoniae* (*Kp*) was not due to *Kp* resistance to fluoroquinolones—*Ef* and *Kp* were both strongly inhibited by all three fluoroquinolones and remained at very low abundance without seeding across all recovery passages (Fig. S5B). *Ef* and *Kp* isolates were also highly susceptible to 20 µM of each fluoroquinolone in monoculture, although *Kp* grew better than *Ef* at 0.1 µM drug concentration (Fig. S5C). The first recovery passage involved drug carryover at a concentration ~0.1 µM, which likely allowed *Kp* to outcompete *Ef* after seeding during R1. To test this hypothesis, we treated an *Ef*-*Kp* co-culture with each of the fluoroquinolones and seeded with an equilibrated *Ef*-*Kp* co-culture (~100:1 *Ef*:*Kp* ratio) at 1% relative abundance (v/v) before R1. As predicted, *Kp* ended up at higher abundance than *Ef* after R1 (Fig. 4E). However, in contrast to the persistent blooming of *Kp* throughout R1-6 in seeded SIC-0, the advantage of *Kp* in pairwise co-culture was transient, and *Ef* grew to high abundance by R4 (Fig. 4E). To test whether the expansion of *Ef* in pairwise co-culture was due to the higher *Ef* to *Kp* ratio in treated pairwise co-culture (~10^–2^) compared to in a community (~10^–3^), we mixed *Ef* and *Kp* at different ratios and passaged the mixture. Even with a starting *Ef*:*Kp* ratio of 10^–4^, *Ef* outcompeted *Kp* after 3–4 passages (Fig. S5D). Therefore, despite *Kp*’s growth advantage during fluoroquinolone treatment, other strains in the community likely also play a role in inhibiting *Ef* growth during recovery.

We next tested whether more recovery passages of the six seeded communities with low *Ef* and high *Kp* (three fluoroquinolones, two replicates each) would allow *Ef* to recover. We revived the fluoroquinolone-treated communities by diluting glycerol stocks of their R6 passage into fresh BHI, grew them to saturation, and propagated the resulting communities for six more passages. We refer to these as passages R8–13 (with the revival passage from glycerol stock as passage R7). The seeded communities all started with low *Ef* and high *Kp* at R8. In two of the six communities (both levofloxacin-treated), *Ef* started to increase in abundance during passages R10 and R12, respectively, accompanied by a decrease in *Kp* abundance (Fig. S5E). To query if the recovery of *Ef* was due to the absence of another competing strain or the presence of another strain facilitating *Ef* growth, we compared the abundance of each ASV across the six seeded communities during passages R8–10. Only one of the 124 ASVs, representing *Bacteroides stercoris* (*Bs*), exhibited lower abundance in the levofloxacin-treated communities compared to the gatifloxacin- and moxifloxacin-treated communities (FDR-adjusted *p*=0.007, one-tailed Mann–Whitney U test), and it was present at ~10^–3^ or lower relative abundance in all six communities. To test whether *Bs* could directly inhibit *Ef* growth, we isolated a *Bs* strain, grew it to saturation in BHI supplemented with heme, and assayed *Ef* growth in the supernatant of the saturated *Bs* culture. *Bs* growth by itself yielded high biomass comparable to *Ef* monoculture, yet its supernatant still allowed substantial *Ef* growth (~36% of *Ef* yield in fresh BHI, Fig. S5F). Thus, the low amount of *Bs* in the community is unlikely to fully inhibit *Ef* growth either by nutrient competition or other non-competitive interactions such as toxin production. We also passaged the six unseeded communities (all starting with low *Ef* and low *Kp*), and found that *Ef* recovered to high abundances in two communities during passages R8–9 (one gatifloxacin- and one levofloxacin-treated, Fig. S5G). In the other four cases, although *Ef* remained low, the communities had very different compositions from each other and from the low-*Ef* seeded communities (Fig. S5H), at both the ASV and family levels. For the two unseeded communities in which *Ef* recovered during passages R8–9, we identified a single ASV in the *Lachnospiraceae* family that was differentially abundant (FDR-adjusted *p*=0.01, one-tailed Mann–Whitney U test) during passages R3–6, and it was present at ~10^–3^ or lower relative abundance in all communities. Taken together, we were unable to pinpoint an ASV or family that could explain the overall suppression of *Ef* growth, suggesting that *Ef* may be inhibited by the collective effects of many species in the communities. Another possible explanation was that the *Ef* population in these fluoroquinolone-treated communities are a resistance subpopulation that becomes less competitive to *Kp*. However, when we re-treated the low-*Ef* communities from R6 with the same fluoroquinolones, *Ef* abundance remained low (0.2±0.6% relative abundance, mean±S.D., *n*=12 samples), arguing against resistance.

**Effects of drug treatment are qualitatively conserved across SICs**

We asked whether drug treatment led to conserved compositional changes across SICs by performing a Principal Coordinate Analysis (PCoA) on the communities resulting from treatment of SIC-0, SIC-MD, and SIC-cip. The communities separated by their source along principal coordinate (PCo) 1 (Fig. S6C, left), with SIC-cip-treated communities further away from SIC-0- and SIC-MD-treated communities, consistent with our observation that SIC-cip has more compositional differences from the other two SICs than they did from each other (Fig. 6A, S6A). Meanwhile, the treated communities largely separated by drugs along PCo 2 (Fig. S6C, right), suggesting that each drug exerts some conserved effects across SICs.

Although they clearly differed in composition, the three SICs tested above were all derived from mice colonized with the same human fecal sample. To further test the generality of our screening results, we next determined the effects of drug treatment on eight SICs derived from different human hosts^2^. Compared to SIC-MD and SIC-cip, these SICs harbored even more diverse strains with distinct evolutionary histories. Nonetheless, these SICs (hereafter, SIC-1,2,3,…,8) still exhibited similar patterns of growth inhibition across drug treatments (Fig. S6D; Spearman’s *ρ*≥0.77, *p*<10^–10^). Three drugs (oxytetracycline, rolitetracycline, and tetracycline) exhibited large variations in growth inhibition. Only SIC-1, 2, 3, and 6 exhibited high growth AUC and high *Enterobacteriaceae* abundance across the three tetracycline-family chemicals. Hence, we hypothesized that the variation was related to the susceptibility of the distinct *Enterobacteriaceae* family members^2^ in these SICs (Fig. S6E). We isolated an *Enterobacteriaceae* strain from each of the eight SICs (Methods) and assayed its susceptibility to the three drugs. The isolates from SIC-1, 3, and 6 exhibited strong resistance to all three drugs (relative growth ≥80% of vehicle-treated control), while all other isolates were susceptible (relative growth <10%). SIC-2 contained at least 2 *Enterobacteriaceae* strains at high abundance, and our isolate turned out to be susceptible. These findings indicate that susceptibilities of certain species (particularly abundant ones) can affect overall community growth dynamics during treatment.

Since these eight SICs were derived from different subjects, the same ASV is highly likely to represent evolutionarily distinct strains in different SICs^3^. Nonetheless, virtually all ASVs that are shared between SICs exhibited strong correlations across drug conditions (Fig. S6F; FDR-adjusted *p*≤0.02). The only exception was the *E. fergusonii* ASV in SIC-6 (FDR-adjusted *p*=0.4), likely because SIC-6 contains another *Enterobacteriaceae* ASV (a *Proteus* species) at much higher abundance (~70% relative abundance compared to ~10% relative abundance of *Ef* in the untreated SIC-6), which presumably affects *Ef* growth.

Taken together, even though SICs derived from different hosts are composed of distinct bacterial strains, they exhibited similar patterns of growth and compositional changes after drug treatment except when high-abundance species exhibited variable resistance across strains.

**SUPPLEMENTAL FIGURES**

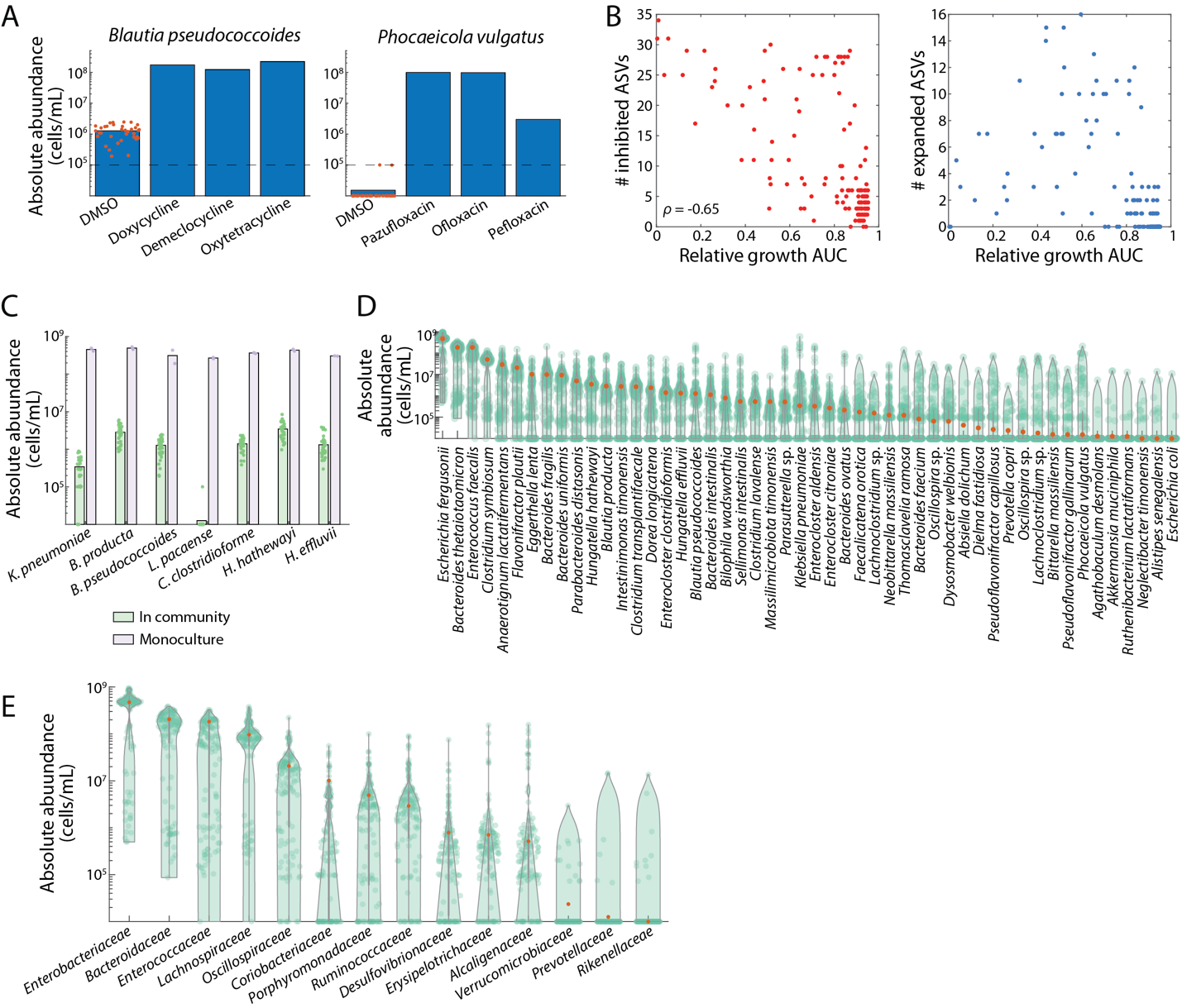

**Figure S1: Drug treatment leads to large-scale abundance changes.** A) Examples of two ASVs that increase in absolute abundance after drug treatment. Red dots: replicates of vehicle-treated control. Dashed line: limit of detection corresponding to relative abundance of 10^-4^. B) Left: larger growth inhibition is correlated with more ASVs exhibiting growth inhibition. Right: intermediate growth inhibition allows more ASVs to expand in abundance. C) Many of the ASVs present at 1% or lower relative abundance in SIC-0 are able to grow to much higher yields (equivalent to >20% of the total yield of the untreated SIC) during monoculture growth in BHI. Dots are biological replicates, *n*=35 for community-level data, and *n*=2 for monocultures. Bars represent the mean across replicates. D,E) Regardless of their initial abundance in SIC-0 (red dots), the abundance of each ASV (D) or family (E) varied >1,000-fold across drug conditions.

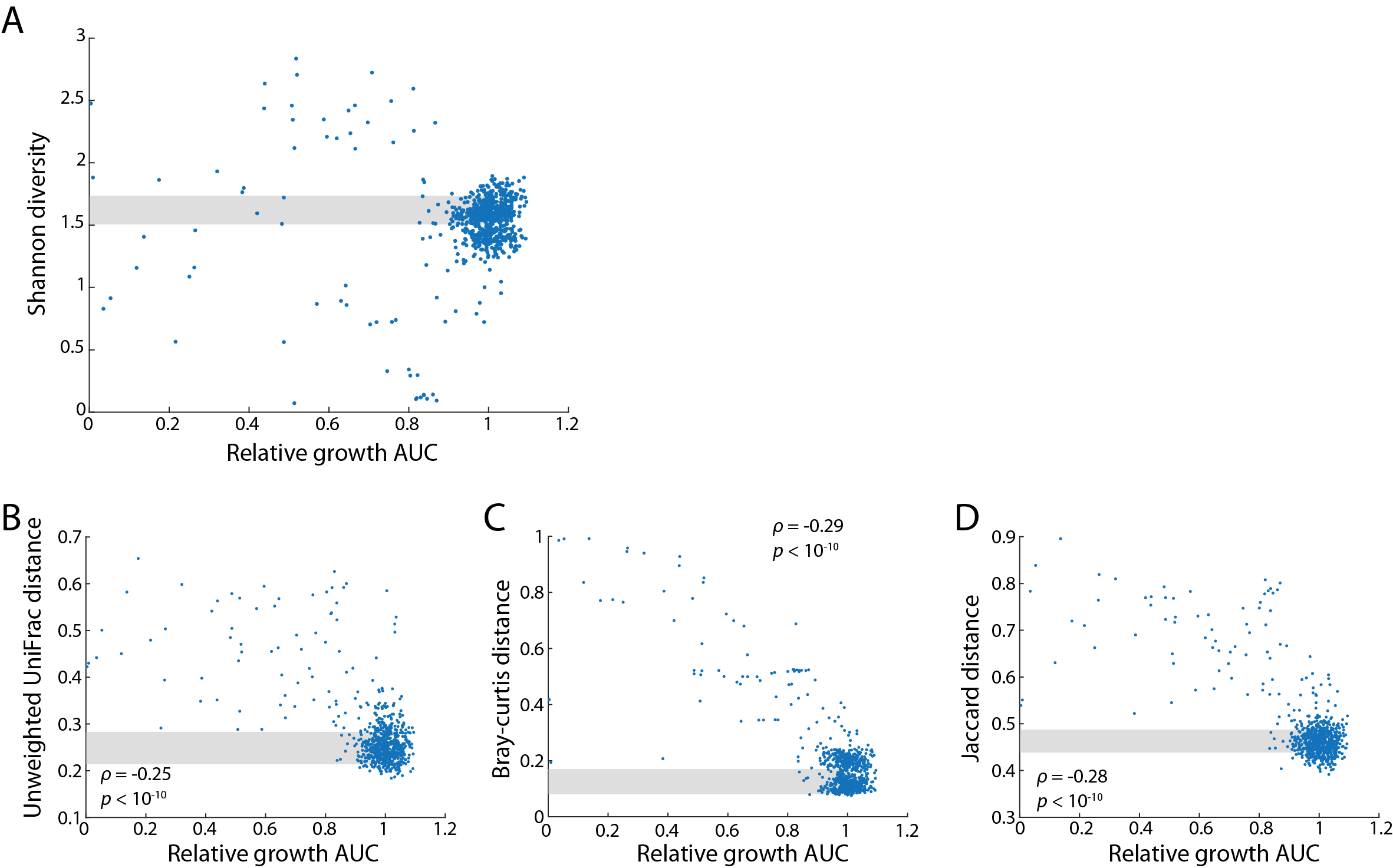

**Figure S2: Growth inhibition in a community is correlated with compositional changes.** A) There was no systematic trend between degree of growth inhibition and alpha diversity as measured by Shannon diversity. B–D) Across treated communities, relative growth AUC was negatively correlated with compositional distance as measured by unweighted UniFrac (B), Bray–Curtis (C), and Jaccard (D) distance to the vehicle-treated control. Shaded areas represent the mean±1 S.D. for vehicle-treated controls.
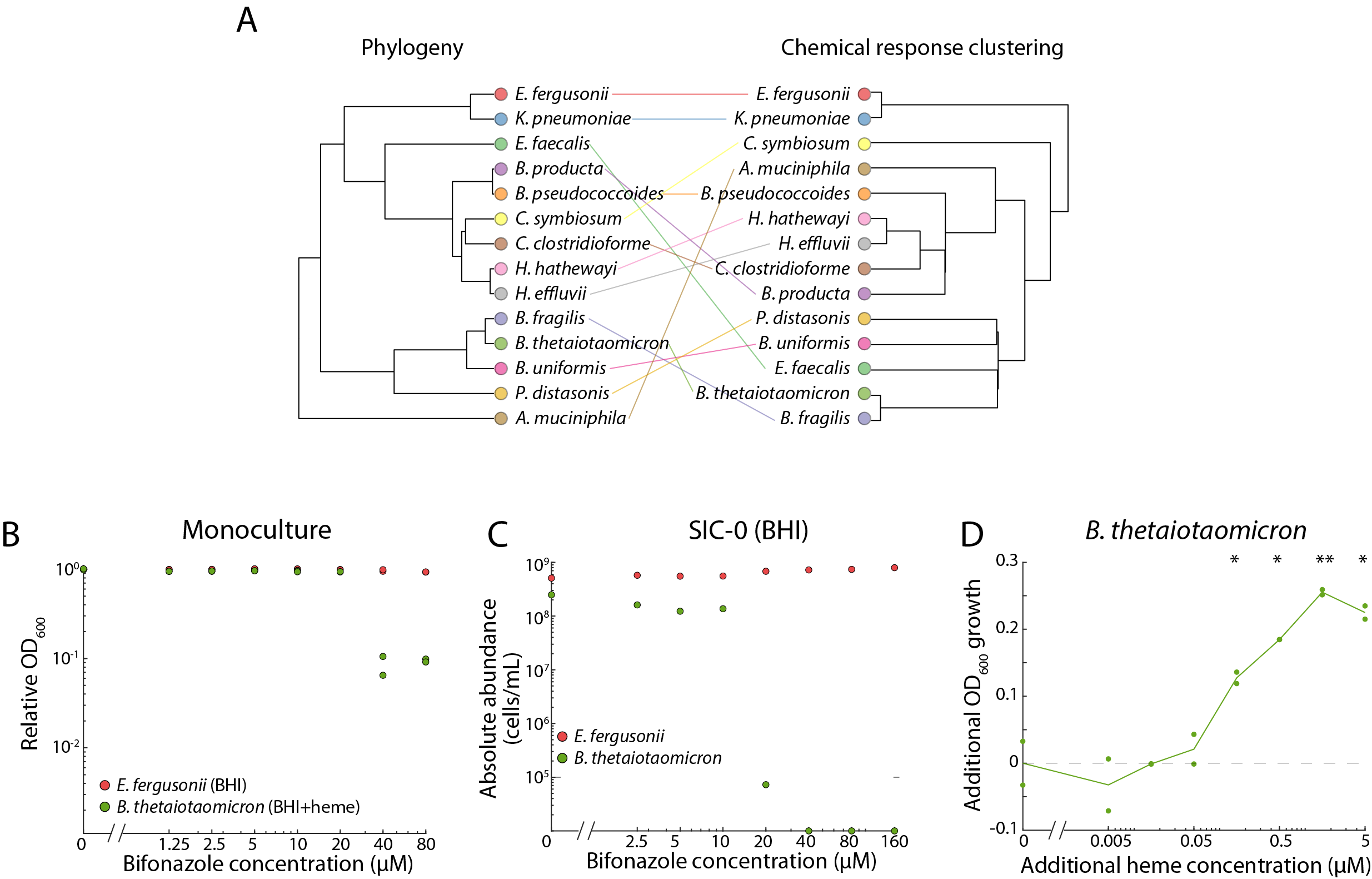

**Figure S3: Growth inhibition in a community is related to inter-species interactions.** A) Phylogeny of bacterial isolates (left) juxtaposed with their growth responses to 20 µM drug treatments (right). Phylogenic distances were calculated from multiple sequence alignments of the V4 region in the 16S rRNA gene. Chemical response distances were calculated as Euclidean distances of their relative growth yield across 43 chemicals. Lines connect the same species to aid visualization. Overall, clustering based on phylogeny and chemical responses resulted in distinct outcomes (Mantel statistic *r*=0.38). B) *B. thetaiotaomicron* (*Bt*) growth in monoculture was not impacted by bifonazole treatment at concentrations 20 µM or lower. C) In a community, *Bt* was strongly inhibited by 20 µM bifonazole. D) At concentrations 0.16 µM or higher, supplemented heme promoted *Bt* growth in BHI. Dashed line is *Bt* growth without additional heme. Data are mean of two replicates (shown as individual dots). *: *p*<0.05, **: *p*<0.01, one-tailed Student’s *t*-test.

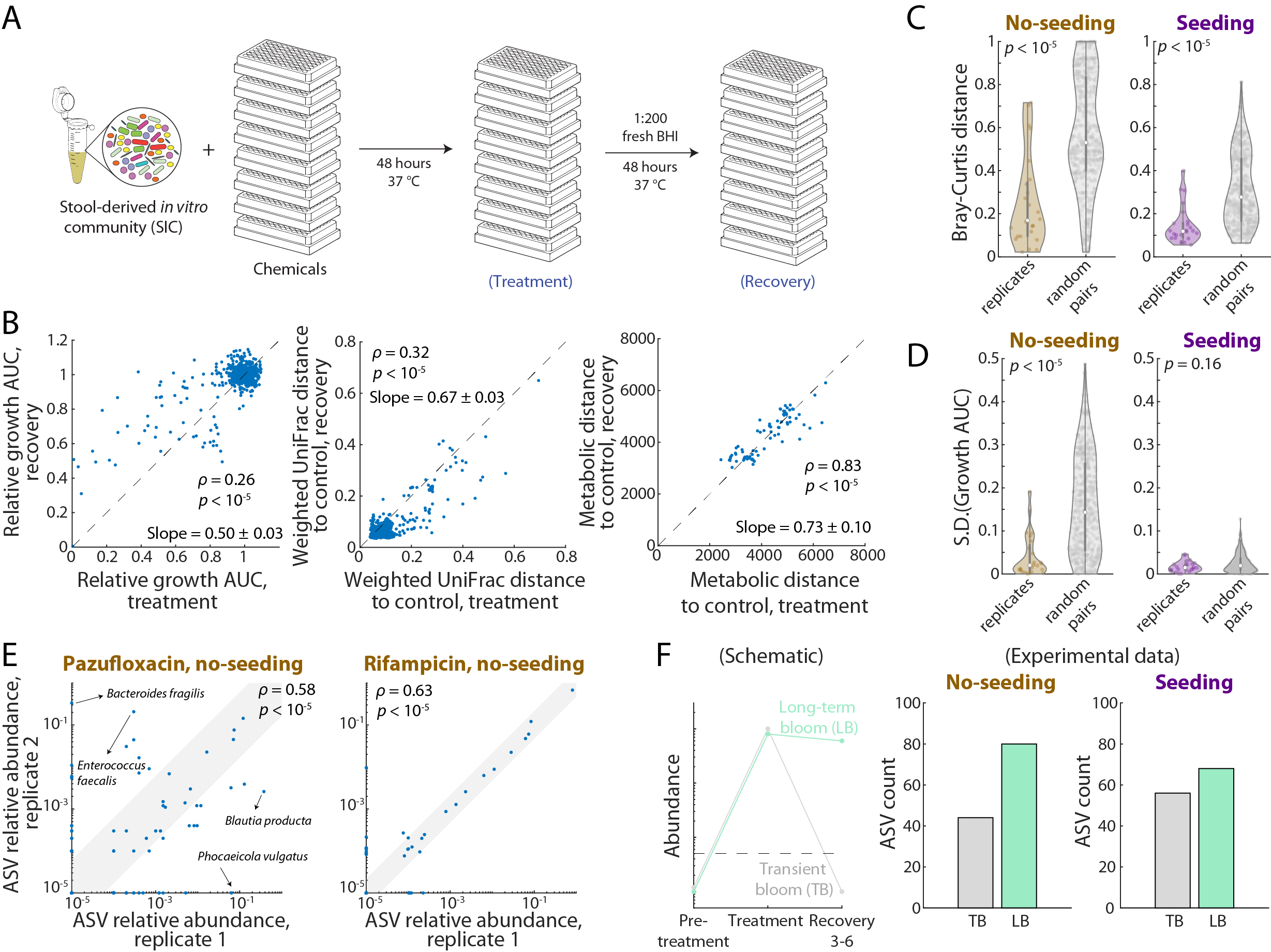

**Figure S4: Community recovery from drug treatment is hampered by extinction of some species.** A) Schematic of the recovery passage after the initial screen in Fig. 1B. B) A single passage after treatment enabled partial recovery of community growth (left), composition (middle), and metabolomic profile (right). Dashed lines: *y*=*x*. C,D) After multiple recovery passages (Fig. 4A), the replicate pairs exhibited more similar compositions compared to random pairs (C). Growth dynamics were also more similar between replicate pairs without seeding (D, left). With seeding (D, right), all communities grew similarly (Fig. 4B). E) Left: During pazufloxacin treatment, the two replicates exhibited different compositions, particularly involving the abundance of certain high-abundance ASVs; nonetheless, there was still a positive correlation across all ASVs. Right: During rifampicin treatment, replicate community compositions were strongly correlated across all ASVs. Shaded areas represent 95% confidence intervals of linear fits. F) Left: ASVs were defined as transient (TB) or long-term (LB) blooming based on their behavior during recovery passages R3-6. Dashed line: limit of detection. Middle and right: Regardless of seeding condition, many ASVs exhibited long-term blooming after drug treatment.

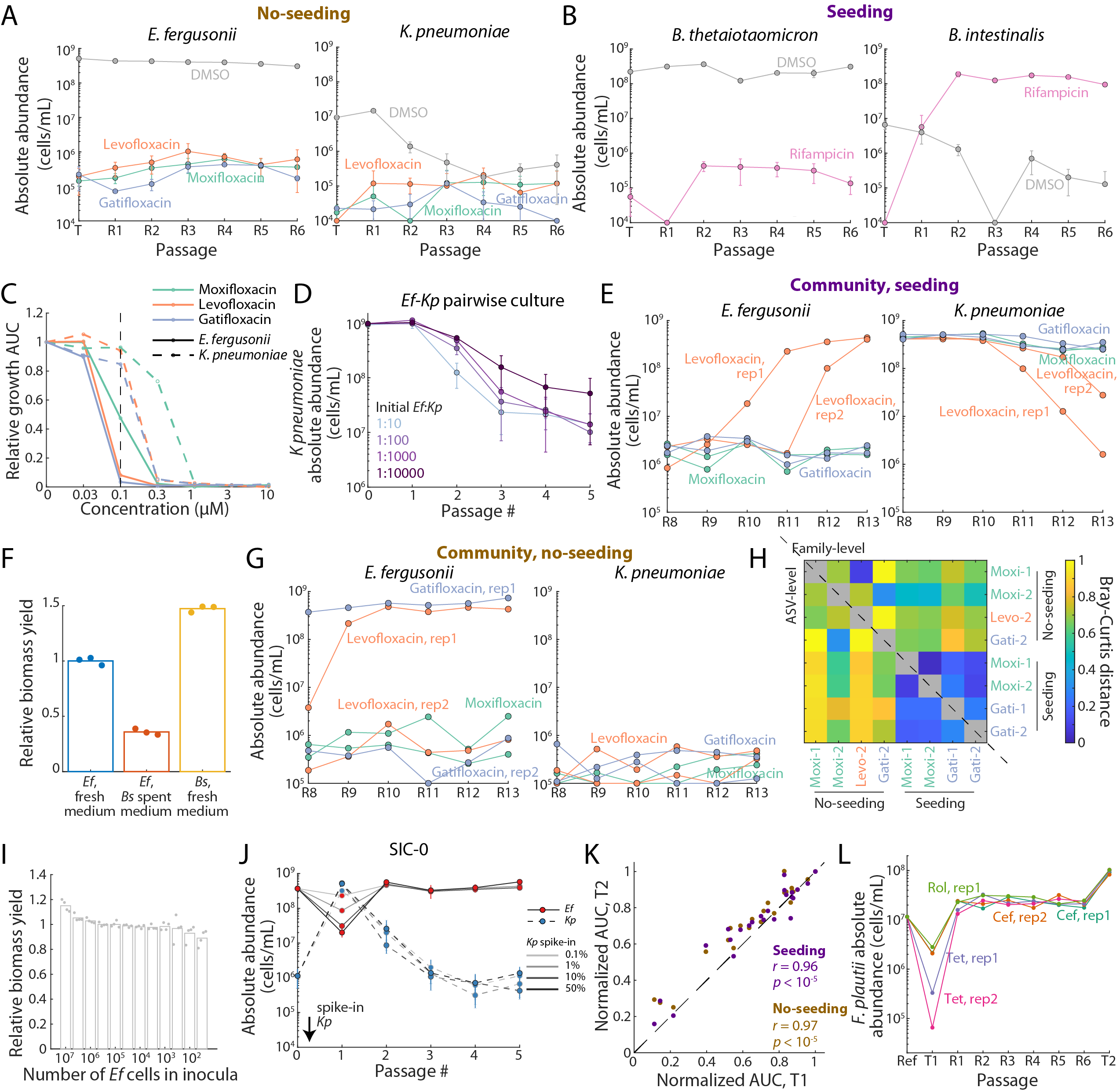

**Figure S5: Strain swapping in drug-treated communities.** A) Without seeding, fluoroquinolone treatment strongly suppressed both *E. fergusonii* (*Ef*) and *K. pneumoniae* (*Kp*). Data are mean±1 S.D. across *n*=2 replicates. B) Rifampicin treatment led to a strain swap in which *B. thetaiotaomicron* was replaced by *B. intestinalis*. C) *Kp* is more resistant than *Ef* to fluoroquinolones, with much higher relative growth AUC at 0.1 µM concentration. D) Regardless of the initial mixture ratio between *Kp* and *Ef*, *Kp* abundance decreased by 10- to 100-fold in a pairwise co-culture over five passages. Data are mean±1 S.D. across *n*=3 replicates. E) When the six seeded communities shown in Fig. 4D were passaged further, *Ef* eventually recovered in some of them. F) Spent medium from *B. stercoris* monoculture still allowed for *Ef* growth. G) When the no-seeding communities in (A) were passaged further, *Ef* recovered in some of them. H) Across all communities (both no-seeding and seeding) with low *Ef* after R13, they exhibited highly different compositions at the ASV and family levels as measured by Bray-Curtis distance. I) In monoculture, *Ef* growth yield remained high regardless of its initial inoculum size. J) SIC-0 was resistant to challenge with *Kp*, always recovered to a low-*Kp*, high-*Ef* state regardless of the initial ratio of *Kp* spike-in. K) Community growth AUC was largely the same between the first and second treatments. *r*: Pearson’s linear correlation; *p*-values are from two-tailed Student’s *t*-tests with *n*=24 for each condition. K) With seeding, *F. plautii* exhibited resistance selection under similar drug conditions as in the unseeded communities (Fig. 5D).

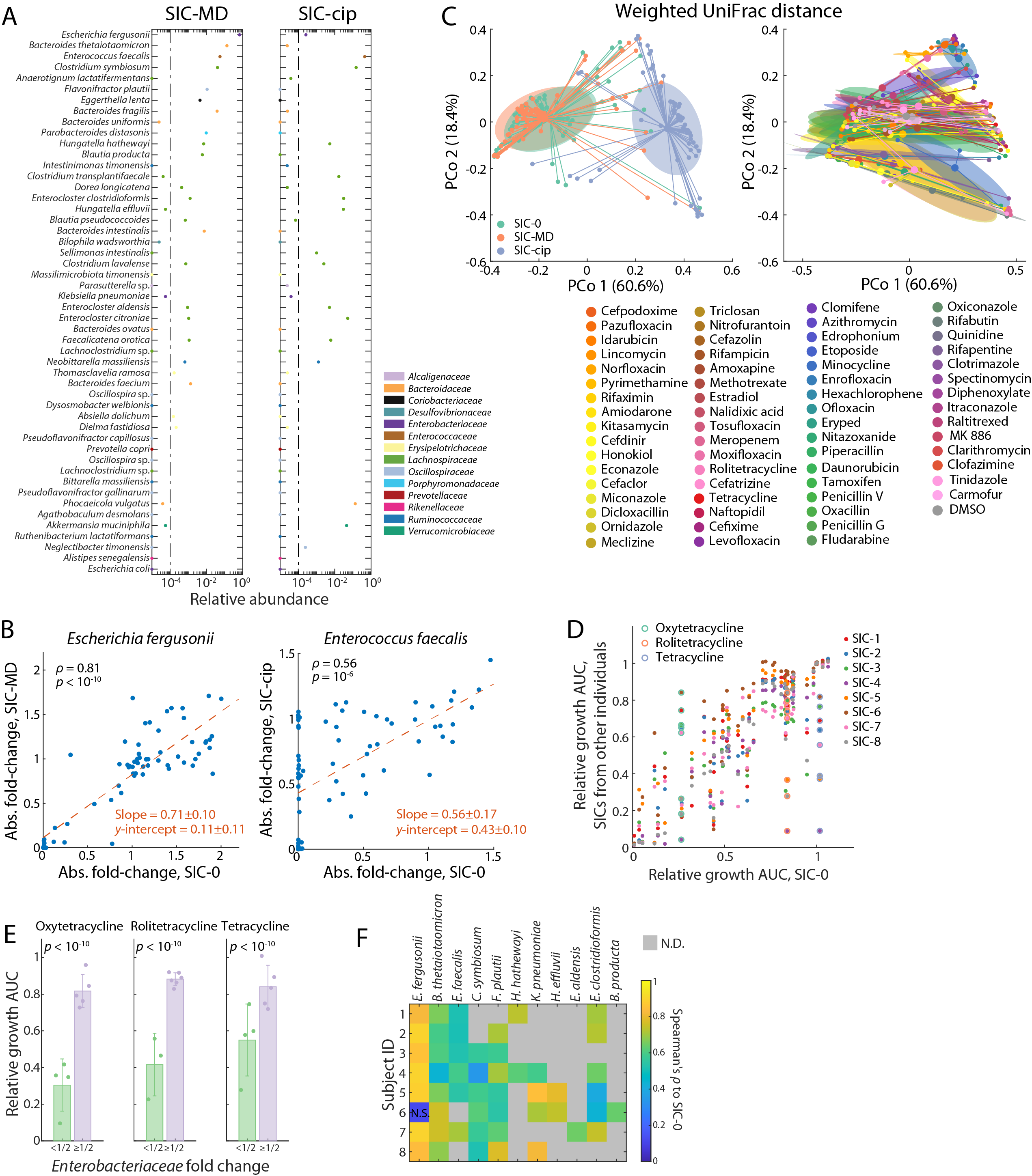

**Figure S6: Effects of drug treatment are largely conserved across SICs.** A) Relative abundances of ASVs detected in both SIC-MD and SIC-cip. The ASVs are sorted by their abundance in SIC-0 (Fig. 1C). Each ASV is colored by its family. These SICs contain some of the same ASVs as SIC-0, but the relative abundances of many overlapping ASVs differ across SICs. B) Absolute fold-change of *E. fergusonii* (left) and *E. faecalis* (right) ASVs across SICs. In both cases, the fold changes in two SICs were correlated but deviated from the line *y*=*x*. C) PCoA plots of the communities resulting from drug treatment of SIC-0, SIC-MD, and SIC-cip. The pre-treatment SIC (left) and drug condition (right) both affected community composition. Small dots are colored by pre-treatment SIC (left) or drug condition (right), and large dots are the corresponding centroids. Ellipses represent the 60% confidence intervals of the dots. D) SICs derived from other human subjects exhibited similar growth dynamics as SIC-0, although there were large variations for some tetracycline-group chemicals. E) The variations in growth AUC for tetracycline-group chemicals can be largely explained by the susceptibility of *Enterobacteriaceae* ASVs in each SIC, such that *Enterobacteriaceae* susceptibility is correlated with large growth inhibition. >(<)1/2: the corresponding changes in *Enterobacteriaceae* family under drug treatment is more (or less) than 1/2. *p*-values are from two-tailed Student’s *t*-tests. F) For all ASVs shared between SIC-1-8 and SIC-0, we observed strong positive correlations in abundance change across all drugs, with the exception of the *E. fergusonii* ASV in SIC-6. N.D.: no data (due to the absence of the ASV in one or both SICs). N.S.: not significant.

**SUPPLEMENTAL TABLES**

**Table S1: List of NCATS drugs and their primary targets.**

**Table S2: List of drugs selected for monoculture susceptibility testing.**

**SUPPLEMENTAL REFERENCES**

1	Treffers, H. P. The linear representation of dosage-response curves in microbial-antibiotic assays. *Journal of bacteriology* **72**, 108-114 (1956).

2	Aranda-Diaz, A. *et al.* Assembly of gut-derived bacterial communities follows" early-bird" resource utilization dynamics. *bioRxiv*, 2023.2001. 2013.523996 (2023).

3	Roodgar, M. *et al.* Longitudinal linked-read sequencing reveals ecological and evolutionary responses of a human gut microbiome during antibiotic treatment. *Genome research* **31**, 1433-1446 (2021).
